# High conservation of the dental plaque microbiome community across populations with differing subsistence strategies and levels of market integration

**DOI:** 10.1101/2022.07.27.501666

**Authors:** Irina M. Velsko, Sandrine Gallois, Raphaela Stahl, Amanda G. Henry, Christina Warinner

**Affiliations:** Department of Archaeogenetics, Max Planck Institute for Evolutionary Anthropology, Leipzig, Germany 07745; Faculty of Archaeology, Leiden University, Leiden, the Netherlands 2333 CC; Faculty of Biological Sciences, Friedrich Schiller University, Jena, Germany 07743; Department of Anthropology, Harvard University, Boston, MA, USA 02138

**Keywords:** Oral microbiome, dental plaque, diet, Baka, Nzime, industrialization, subsistence

## Abstract

Industrialization - including urbanization, participation in the global food chain, and consumption of heavily processed foods - is thought to drive substantial shifts in the human microbiome. While diet strongly influences stool microbiome composition, the influence of diet on the oral microbiome, and particularly dental plaque, is largely speculative. Here we investigated whether dental plaque microbial communities are distinctly different across populations with dissimilar subsistence strategies and degree of industrialized market integration. Using a metagenomic approach, we compared the dental plaque microbiomes of Baka foragers and Nzime subsistence agriculturalists in Cameroon (n = 46) with the dental plaque and calculus microbiomes of highly industrialized populations in North America and Europe. We found that differences in microbial taxonomic composition between populations were minimal, with high conservation of abundant microbial taxa and no significant differences in microbial diversity related to dietary practices. Instead, we find that the major source of variation in dental plaque microbial species composition is related to tooth location and oxygen availability, and may be influenced by toothbrushing or other dental hygiene measures. Our results support that dental plaque, in contrast to the stool microbiome, maintains an inherent stability against ecological perturbations in the oral environment.

## Introduction

The human mouth is a complex environment, containing up to seven distinct surfaces that host distinct microbiomes ^1,2^. Dental plaque, which forms on tooth surfaces, is a dense biofilm containing predominantly bacteria, the species composition of which is strongly linked to oral and systemic health and disease ^3,4^. The dental plaque microbial community has been relatively well-characterized for individuals in heavily industrialized countries, particularly in North America and Europe, where heavily processed diets are consumed ^2,5^. However, little is known about the structure and function of microbial dental plaque communities among individuals outside of industrialized systems and in association with more traditional subsistence strategies.

As microbiome research has expanded to include more diverse human groups, increased attention has been paid to the role of industrialization in influencing the human microbiome. For example, the stool microbiome of industrialized and non-industrialized societies consistently shows distinct compositional differences, with stool from non-industrialized groups being characterized by higher microbial diversity, as well as higher proportions of *Prevotella* ^6–12^ and the presence of treponemes ^6,8–11^, eukaryotic intestinal parasites ^9,11^, and unique bacteriophages ^13^. Such distinctions have largely been linked to dietary and subsistence differences, as well as lifestyle changes with respect to sanitation, antibiotic exposure, and urbanization ^6,7,9,11,14,15^. The clear difference in stool sample taxonomic composition between industrialized and non-industrialized populations is often extrapolated to represent a generalized shift in the human microbiome associated with industrialization, yet similarly consistent differences have not been reported in oral microbiome studies.

Of the studies to date that have investigated the role of diet on oral microbiome composition, simple sugar intake has been associated with microbial shifts ^16^, but broader microbial changes related to differences in subsistence or urbanization, industrialization, and/or hygiene are subtle and often inconsistent ^17^. For example, for hunter-gatherers in the Amazon basin, the Philippines, and Central Africa, within-sample diversity of oral bacteria in saliva and oral mucosa has been reported to be both similar to those of industrialized populations ^18,19^ and higher than those of subsistence farmers or more urbanized populations living in close geographic locations ^14,20,21^.

Likewise, taxonomic composition and between-sample microbial diversity of these populations have been reported to differ from industrialized populations ^18,21^, but also to be highly similar ^14,19,20^. Although these contradictory findings may be related to a variety of study-specific factors (e.g., differences in study design, sample collection and preparation, and data processing and analysis), similar methodological heterogeneity also characterizes gut microbiome studies, which in contrast have yielded consistent findings. Terminology describing the groups under study may be another factor in the inconsistencies of reported findings. The dichotomy of industrialized/non-industrialized populations is a mis-representation of continuous scales of integration into the market economy, consumption of industrially-processed foods, and level of urbanization ^22^. In particular, the term non-industrial is broadly used to refer to groups with substantial heterogeneity in their placement across these scales, and therefore, across numerous studies, populations described as non-industrial cannot be equated ^22^.

One study-design issue that may be preventing oral microbiome studies from detecting consistent differences is the variety of sampling sites in the mouth. Saliva, oral mucosa, tongue, and dental plaque harbor distinct microbial communities, and the species profiles of these different sites cannot be directly compared ^1,2,23,24^. Saliva, which is easy and non-invasive to collect, is not a true microbiome, but rather a dynamic assemblage of microbes shed from other oral sites, especially the tongue, oral mucosa, and throat ^25–27^. Therefore, saliva studies may be more prone to stochastic results than studies using solid sites that allow bacterial adherence, such as dental plaque or keratinized gingiva. Dental plaque, a dense and highly structured biofilm, is relatively well-characterized in North American and European populations ^4,5^, and population-level comparisons of dental plaque rather than saliva may lead to more consistent results related to the impact of integration into the global food chain, urbanization, and industrialization on oral bacteria. In addition, calcified dental plaque preserves well in the archaeological record as dental calculus ^28,29^, potentially allowing long-term inferences to be made about factors influencing the evolution, ecology, and function of the oral microbiome ^30^.

Here we investigate the dental plaque composition of individuals from two ethnic groups in Cameroon whose livelihoods are based on foraging and small-scale farming, the Baka and the Nzime, in order to better understand oral microbiome diversity within these groups. While both groups practice varying degrees of foraging and small-scale farming, Baka are historically more nomadic, depending more on foraging, and incorporate more wild and foraged foods into their diet, while Nzime are more sedentary, relying mostly on agriculture, and incorporate more domesticated and cultivated foods into their diet. Nevertheless, despite these differences in subsistence and lifestyle, we find that the dental plaque microbial communities of Baka and Nzime individuals (hereafter BN) are highly similar to those of both dental plaque and dental calculus from heavily industrialized populations in the US and Europe (hereafter USE). Further, no associations between dental plaque taxonomic composition and subsistence strategy were identified. These findings are consistent with recent archaeogenetics studies showing the long-term stability of dental calculus microbial communities within diverse human populations extending back to the late Pleistocene ^30,31^, as well as experimental studies examining dental plaque changes over short and medium time frames ^32,33^.

Collectively, these findings support a characteristic stability of oral microbiota that is distinct from the gut ^34^ and suggest that the human microbiome does not undergo monolithic changes in response to major social, economic, and lifestyle changes. Rather, each body site is affected and constrained by distinct factors. Characterizing the diversity and evolutionary history of the oral microbiome may be key to both understanding and treating microbially-mediated diseases of the oral cavity.

## Results

### Subsistence and lifestyle in southeastern Cameroon

This study was conducted as part of a broader investigation of foraging strategies among the Baka people of southeastern Cameroon, with a specific focus on the importance of plant dietary resources in forested tropical environments. We partnered with neighboring members of two distinct ethnic groups, the Baka, a Ubangian speaking people who practice foraging and small-scale cultivation, and the Nzime, a Bantu-speaking people who practice subsistence-level agriculture ^35,36^, in order to better understand the influence of diet and lifestyle on oral microbial communities. Although living in the same area (Figure 1), Baka and Nzime lifeways greatly differ.

**Figure 1.**
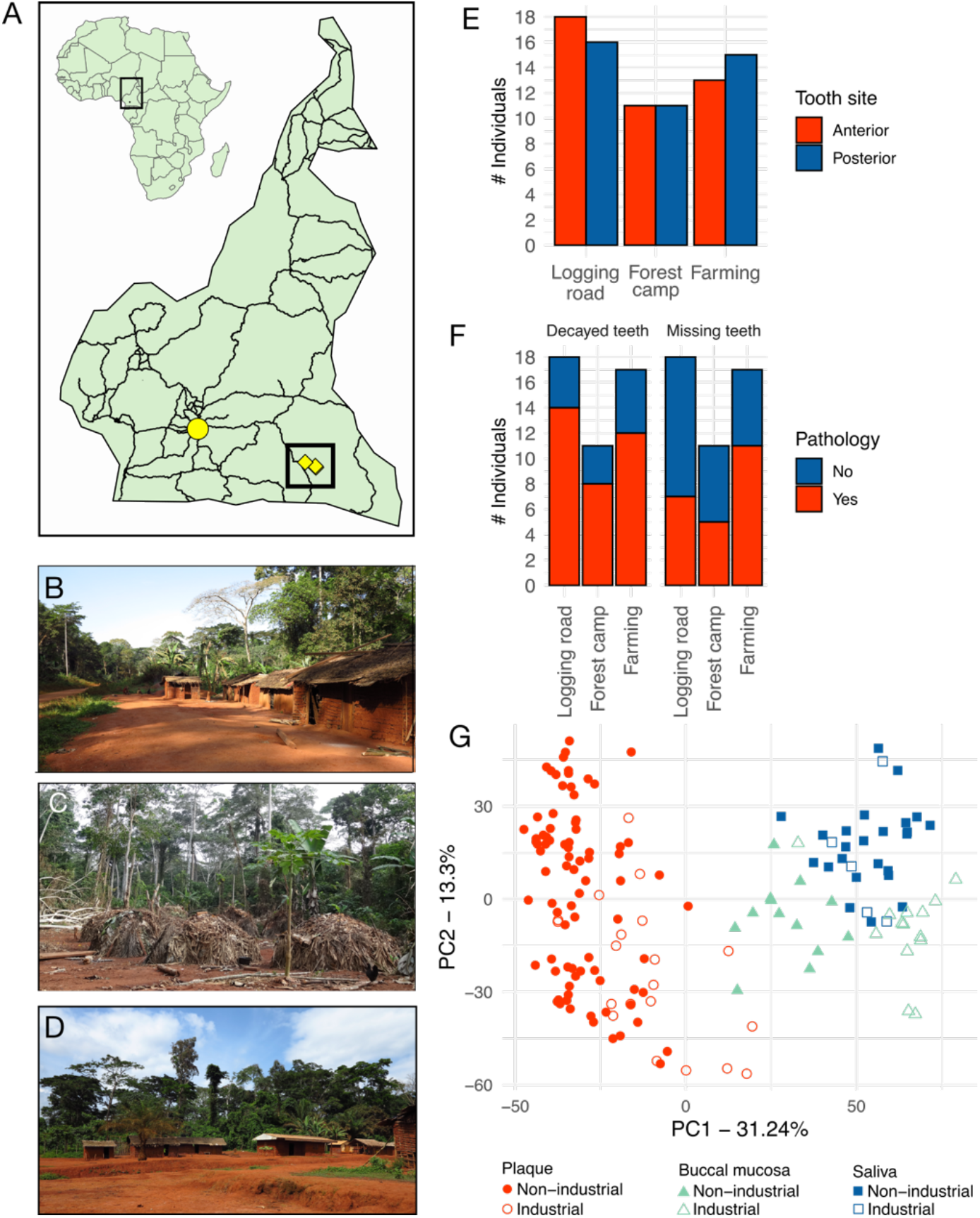
Sample collection locations. **A**. Location of the villages included in this study. The inset of Cameroon shows the major roadways, and the capital Yaounde marked as a yellow circle. The locations of the villages are indicated by yellow diamonds in the highlighted box. The villages are in close proximity and the diamonds overlap.**B**. Baka village along a logging road. **C**. Baka forest camp village. **D**. Nzime village along a logging road. **E**. Number of individuals with missing or decayed teeth. No significant differences were found between groups. **F**. Number of individuals with anterior and posterior samples. No significant differences were found between groups. **G**. PCA plot of plaque, buccal mucosa, and saliva microbial composition from groups with high market integration (industrial) and low market integration (non-industrial). Photo credits: Sandrine Gallois.

The Nzime rely on subsistence agriculture, cultivating mostly cassava and plantains as major crops, and also producing cacao for trade. The Baka rely on a more mixed subsistence strategy, combining foraging (hunting, gathering, and fishing) with small-scale farming ^37^. The Baka and Nzime have long had an economic relationship based on the exchange of foraged forest and farmed agricultural products. The diets of both the Baka and the Nzime are primarily plant-based, with a focus on cassava and plantains, supplemented by bushmeat and cultivated or wild vegetables. While the main components of Baka and Nzime diets overlap (Supplemental table S2), Baka diets have a higher degree of dietary seasonality and they consume a wider diversity of wild products in their daily meals, although such diversity tends to decrease with higher integration into the market economy ^37,38^.

Both the Baka and Nzime have limited access to market economy goods through small local shops and traders traveling along logging roads, but the Nzime, due to their higher income from agricultural surplus, have greater access to market goods and integrate a higher proportion of processed foods obtained from local shops into their diet. Toothpaste is not typically available, and dental cleaning is uncommon, particularly among the Baka. The Nzime participants were from 2 farming villages situated along logging roads. The Baka participants were from 3 villages, two of which were situated along logging roads, and one of which was a forest camp. Individuals living in the forest camp have much lower consumption of market economy goods than those living along the logging roads. Therefore, for analysis we grouped the plaque samples by village location (logging road and forest camp for Baka, and farming for Nzime) (Figure 1A-D).

Baka and Nzime volunteers in this study participated under informed consent. Before the onset of the research we received the Free Prior Informed Consent from both the local authorities of the villages and all the volunteers. The study received the research permit from the Ministry of Scientific Research and Innovation (MINRESI n°000115/MINRESI/B00/C00/C10/C12 and n°000116/MINRESI/B00/C00/C10/C12). The study design was approved by the Ethical Committee from the Ministry of Health of Cameroon (n°2018/06/1049/CE/CNERSH/SP) and the ethics committee of Leipzig University (196-16/ek). It also adheres to the Code of Ethics of the International Society of Ethnobiology ^39^. As part of the study, participants received a dental health assessment and dental cleaning, and the results of the study were returned to the participants at the conclusion of the study.

### Generation of metagenomic data from Baka and Nzime dental plaque

Where possible, we collected from each participant two dental plaque samples, one from the anterior teeth (incisors, canines) and the other from the posterior teeth (premolars, molars). This sampling strategy was adopted because previous studies have found small but significant differences between microbial communities on anterior and posterior teeth related to oxygen tolerance. From 46 individuals, we successfully built 85 shotgun DNA libraries: 57 libraries from 29 Baka individuals (27 paired anterior and posterior samples, 2 unpaired samples, and 1 additional anterior sample from a paired set), and 28 libraries from 17 Nzime individuals (11 paired anterior and posterior samples and 6 unpaired samples) (Figure 1E, Supplemental table S1). The average sequencing depth across samples was 14.5 ± 3 million paired-end reads, which is sufficient to obtain an accurate species profile ^40^. Samples were taxonomically profiled using MALT and a database of bacteria and archaea from NCBI RefSeq. Eighty-four of 85 libraries contained predominantly known oral bacteria, indicating that they were not subject to overgrowth of environmental taxa during storage and shipping (Supplemental figure S5); one library was built from insufficient biological material and was removed for resembling blanks (see Methods). No significant differences were observed in diversity between extraction or library batches (Figure S6).

### Oral health comparison of Baka and Nzime individuals

Dental health is known to influence dental plaque species composition, such that differences in dental health between comparison groups can confound the results of comparative microbiological studies by obscuring differences stemming from other factors. Therefore, we compared the dental health of the Baka and Nzime participants in this study. We found that dental health was not substantially different between the Baka and Nzime with respect to the number of decayed teeth, missing teeth (Figure 1F), or teeth with exposed roots (p > 0.05) (Supplemental figures S2, S3). Moreover, dental plaque species composition did not cluster samples based on these measures (Supplemental figure S4).

### Comparison of oral site communities across populations with differing levels of market integration

While microbial profiles have been shown to differ across oral sites in North American and European populations, this has not yet been investigated in more diverse global populations. Several oral sites, such as buccal mucosa and saliva, have been individually studied in groups with differing levels of market integration ^14,18–21^; however, to our knowledge, this is the first microbiome investigation of dental plaque in populations practicing foraging or small-scale agriculture. We tested whether microbiome communities are distinctive between three oral sites - dental plaque, saliva and buccal mucosa - by performing PCA on samples from groups with high and low levels of market integration. The microbial community structure clustered the samples by oral site rather than by population or market integration status (Figure 1G), indicating that the distinct microbial community structure of samples from different oral sites is a general feature across human populations, and samples from different oral sites cannot be directly compared.

### Microbial composition of Baka and Nzime dental plaque

To determine the composition and similarity of microbial communities in Baka and Nzime dental plaque compared to dental biofilms previously reported from heavily industrialized populations, represented by dental plaque from the Human Microbiome Project ^1^ and dental calculus from urban populations in Spain ^30,41^, we first compared the presence and abundance of the 50 most abundant microbial species in these groups (88 species in total; Figure 2A, Supplementalfigure S8, Supplemental table S3). We included both dental plaque and dental calculus because these dental biofilms have slightly different species profiles, representing early and late stages of biofilm maturation, respectively ^41^. Of the 88 species, 22 were among the top 50 most abundant in all 3 groups, indicating high conservation of the most abundant species. USE calculus and BN dental plaque shared 29 of their top 50 most abundant species, while USE dental plaque and BN dental plaque shared 30 of their top 50 species (Supplemental Table S3). BN dental plaque samples appear to have an intermediate species profile that falls between USE calculus and USE dental plaque, suggesting that the absence of a frequent method to reduce biofilm accumulation (i.e., toothbrushing) may contribute to the formation of a more mature dental plaque biofilm among the Baka and Nzime.

**Figure 2.**
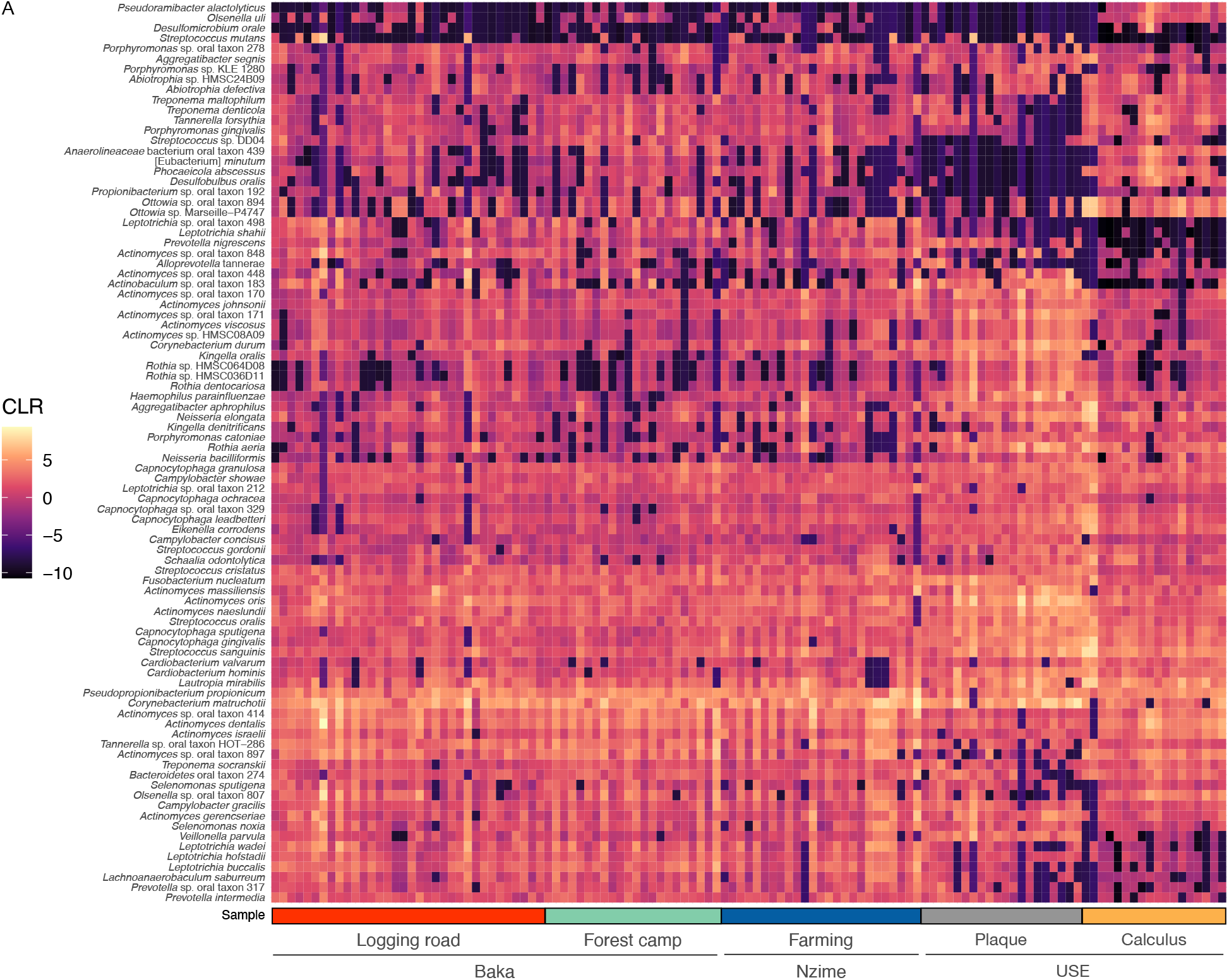
Comparison of abundant oral microbial species across groups. **A**. Heatmap showing the CLR-transformed abundance of the top 50 most abundant species in each group (BN dental plaque, USE calculus, USE dental plaque). Species are hierarchically clustered but samples are not. CLR – centered log ratio.

### Alpha diversity of the Baka and Nzime dental plaque microbial communities

Higher species diversity has been reported in both saliva ^21^ and stool ^6^ samples of populations with lower levels of market integration compared to European and North American populations, so we next looked at whether there is a higher within-sample diversity in dental plaque from Baka and Nzime individuals than in USE dental plaque or calculus. At the species level, we found few significant differences between groups in any metadata tested for the number of taxa or the Shannon Index (Figure 3, Supplemental figure S9, Supplemental table S5). BN dental plaque has on average more species than USE dental plaque and calculus, but the difference is not significant (Figure 3A). We further investigated whether there are differences in the microbial diversity between Baka and Nzime individuals, who differ slightly in their consumption of wild, farmed, and processed foods, and have differences in diet based on gender ^42,43^. The only significant difference in alpha-diversity within these groups was between anterior and posterior teeth (Figure 3B,C). Taken together, microbial diversity in the dental plaque of diverse human groups appears much less affected by dietary and lifestyle differences than in stool.

**Figure 3.**
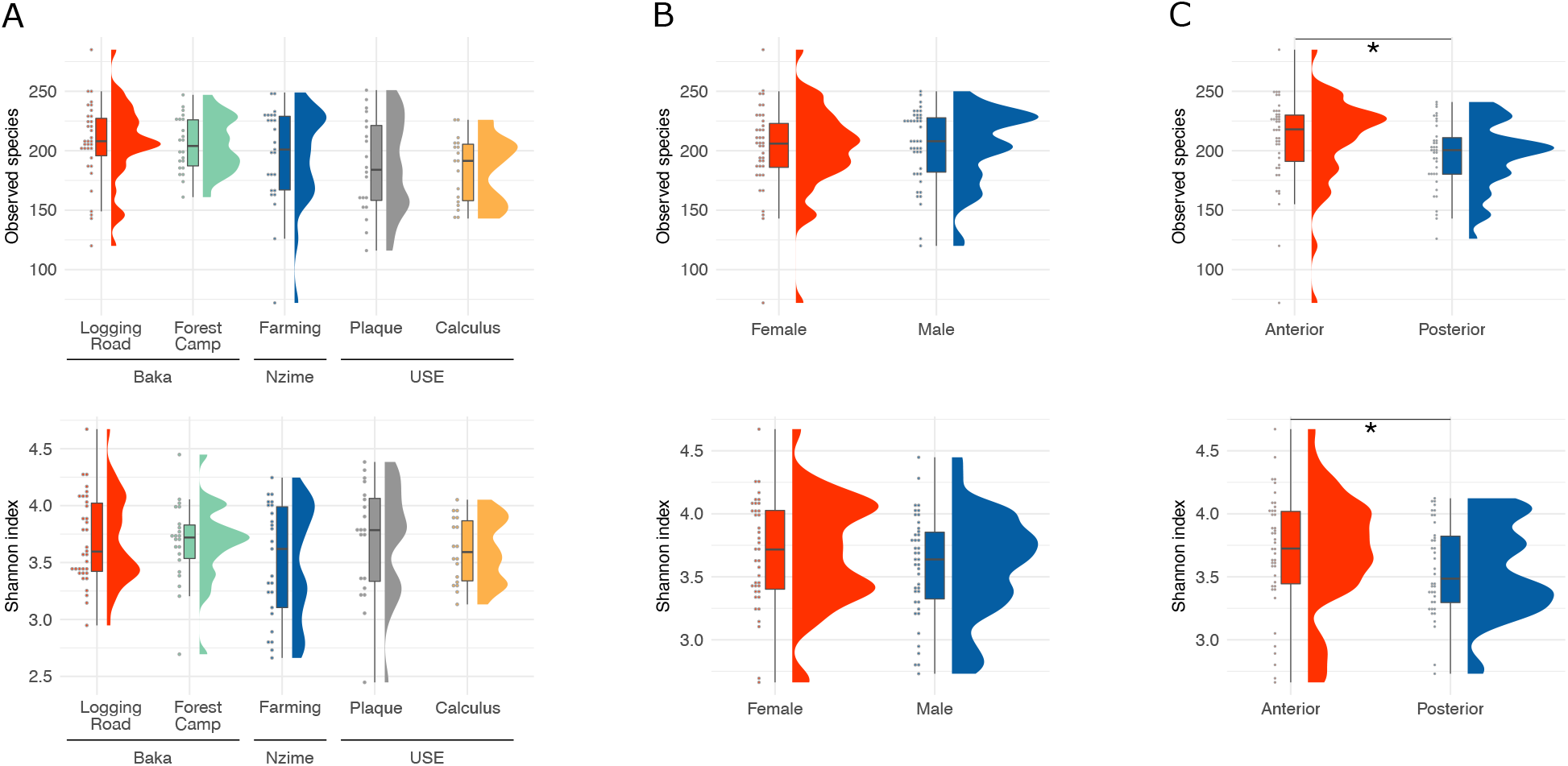
Minimal differences in species diversity exist between Baka, Nzime, and USE oral microbiomes. Alpha-diversity metrics number of species and Shannon index by **A**. market integration and comparison sample source (USE dental plaque and USE calculus). **B**. Baka and Nzime plaque samples only, by gender. **C**. Baka and Nzime plaque samples only, by tooth collection site. * p < 0.05.

### Comparison to diet- and lifestyle-associated changes in the gut microbiome

In previous gut microbiome studies of diverse human populations, notable differences in the most abundant gut taxa were reported between populations with high and low levels of market integration, particularly an inverted abundance of the phyla Bacteroidetes and Firmicutes, and of the genera *Prevotella* and *Bacteroides* within the phylum Bacteroidetes ^6^. To determine whether these differences are also seen in dental biofilms, we calculated the ratio of Bacteroidetes vs. Firmicutes and of *Prevotella* vs. *Bacteroides* within the Baka and Nzime plaque samples, as well as within industrialdental plaque and calculus (Figure 4). The ratio of Bacteroidetes vs. Firmicutes is close to one for all sample groups, with some dispersion in all groups except for USE calculus, in which the samples cluster tightly around one (Figure 4A). No significant differences were detected between groups.

**Figure 4.**
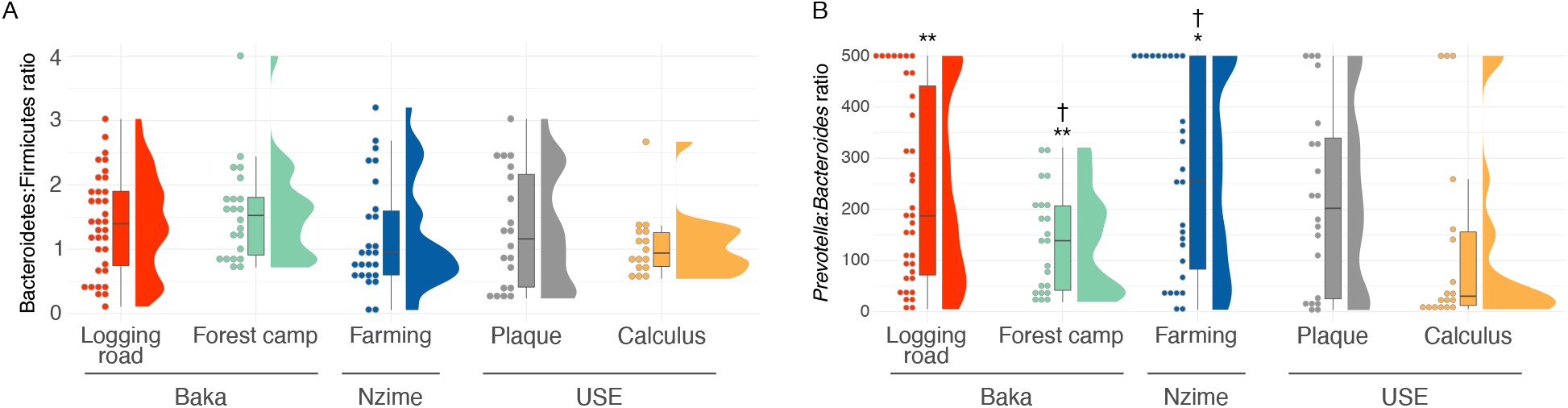
Taxon ratios with distinctive differences in gut microbiomes of populations with high and low levels of market integration. **A**. Ratio of assignments to the phylum Bacteroidetes over assignments to the phylum Firmicutes. **B**. Ratio of assignments to the genus *Prevotella* over assignments to the genus *Bacteroides*. All samples with no *Bacteroides* were assigned a ratio of 500 for plotting purposes, and were excluded from statistical testing. USE - US and European. ** p < 0.01, * p < 0.05 vs Industrial calculus. † p < 0.05 within the group between anterior and posterior samples.

In contrast, the *Prevotella*:*Bacteroides* ratio is much higher than the Bacteroidetes:Firmicutes ratio, averaging above 100 but with substantial dispersion for all sample groups. Again USE calculus, in which the samples cluster more tightly than in the other groups (Figure 4B), is the exception, and all three BN groups are significantly different from USE calculus. The tight clustering of USE calculus samples indicates that calculus profiles are much more similar to each other than the dental plaque samples are to each other, both in USE and BN populations, suggesting that the biofilm climax community observed in calculus is highly consistent across individuals. *Prevotella* are prevalent and abundant in dental plaque, yet few species in *Bacteroides* have been described in dental plaque ^5^, which is reflected in the ratio observed in these samples. Based on these data, it appears that the difference in ratios of Bacteroidetes vs. Firmicutes and *Prevotella* vs. *Bacteroides*, which are clear in gut microbiomes of populations with high and low levels of market integration, is not a characteristic of dental plaque microbiomes.

### *Presence and abundance of* Streptococcus mutans

*Streptococcus mutans* is an oral species strongly associated with dental caries ^44,45^. We observed high counts of *S. mutans* in several BN plaque samples, and investigated the abundance of this species within our dataset related to the number of decayed and missing teeth, both of which are dental pathologies associated with dental caries. *S. mutans* was found in 17 BN plaque samples, 8 USE plaque samples, and no USE calculus samples (Figure 5). Most of the Baka and Nzime individuals with detectable levels of *S. mutans* were over 40 years of age and were female, while all of the USE individuals were under 40 years of age (due to the HMP study inclusion criteria) and were female. In general, for Baka and Nzime individuals with higher proportions of *S. mutans* (Figure 5A), *S. mutans* was the most abundant *Streptococcus* species (Figure 5B), and these individuals had more decayed and missing teeth (Figure 5C), but this was not a consistent pattern. Although *S. mutans* is strongly associated with dental caries, these data support recent studies that have found a highly diverse dental plaque microbiome associated with dental caries, in which *S. mutan*s may not be a central component ^46–49^.

**Figure 5.**
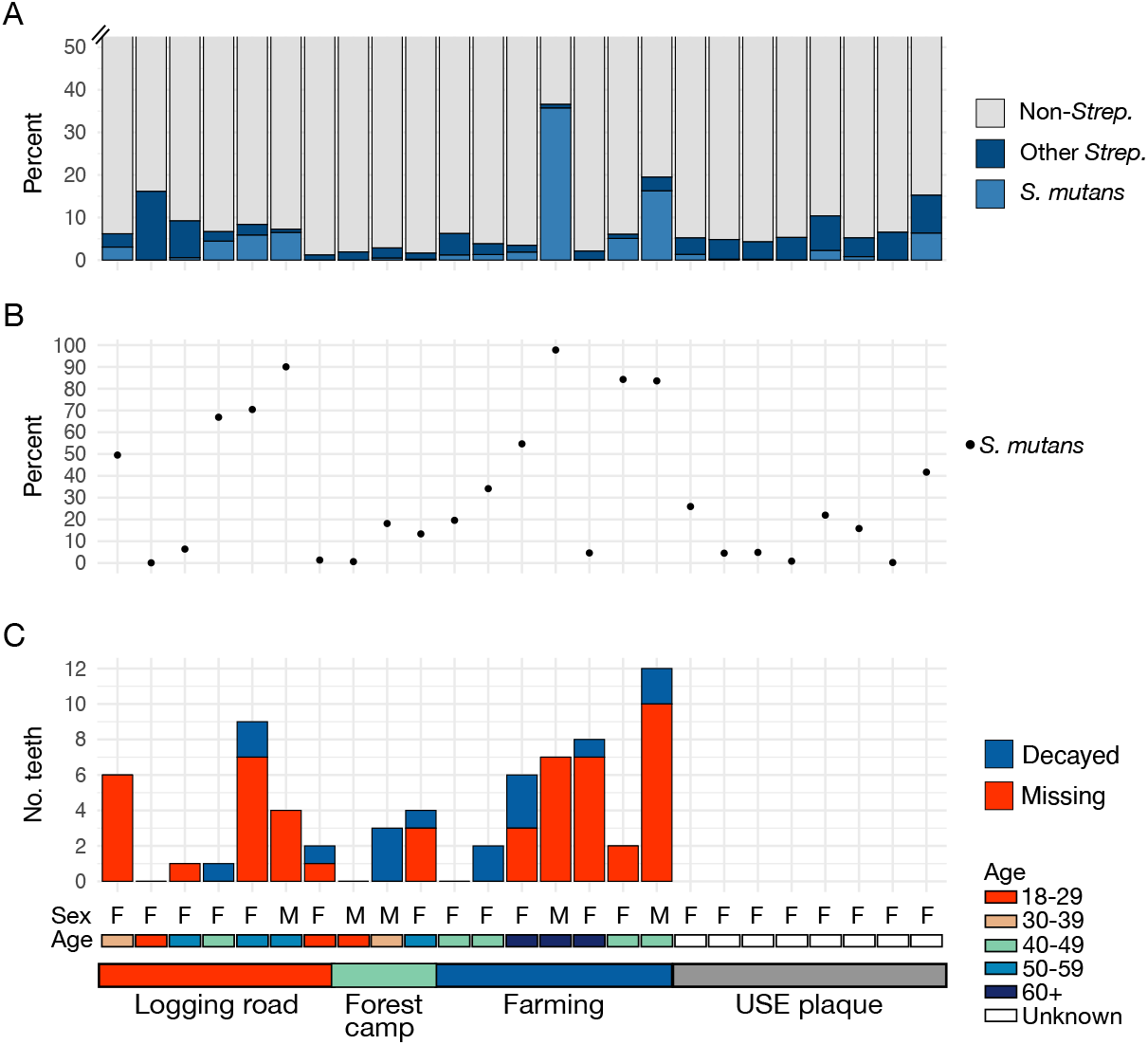
*Streptococcus mutans* proportions in dental plaque. **A**. Percent of reads assigned to all genera except *Streptococcus* (gray), all *Streptococcus* species except *S. mutans* (dark blue), and *S. mutans* (light blue). **B**. Percent of reads assigned to *S. mutans* out of reads assigned to all *Streptococcus* species. **C**. Number of missing and decayed teeth.Data is not available for the USE plaque individuals, but exclusion criteria for the HMP study excluded all individuals with teeth missing for dental disease-related causes and individuals over 40 years old. Only individuals for which *S. mutans* is present are shown.

### Microbial community structure in Baka and Nzime dental plaque

We next asked whether the species diversity profiles of BN dental plaque and USE dental plaque and calculus are distinct, as has been shown for the stool microbiome of groups with different levels of market integration. To address this, we restricted our analysis to the species that contribute most to the variation in our dataset (Supplemental figures S10, S11, Supplemental table S6), as determined by factor analysis, and performed beta-diversity analysis followed by principal components analysis (PCA) (Figure 6, Supplemental Figure S13). Within the PCA plot, USE dental plaque and calculus samples separate along PC2, which is consistent with known species compositional differences between the two sample types ^41^. Baka and Nzime dental plaque diversity falls intermediate between USE dental plaque and calculus samples, falling neither entirely within either group, nor entirely separately (Figure 6A). Of the metadata categories examined (Figure 6, Supplemental figure S13), only tooth site (anterior *vs*. posterior) shows patterning, and this appearsto be driven predominantly by the BN dental plaque samples (Supplemental figure S13D). The BN anterior plaque samples plot closer to USE calculus than posterior plaque, a trend that is maintained when the PCA is repeated using BN dental plaque samples from only a single tooth site (Supplemental Figure S14). This sample clustering suggests that there may be taxa that are more prevalent and/or abundant in the posterior BN plaque than either the anterior BN plaque or the USE calculus samples, which separates them out from the anterior BN and USE samples along PC1.

**Figure 6.**
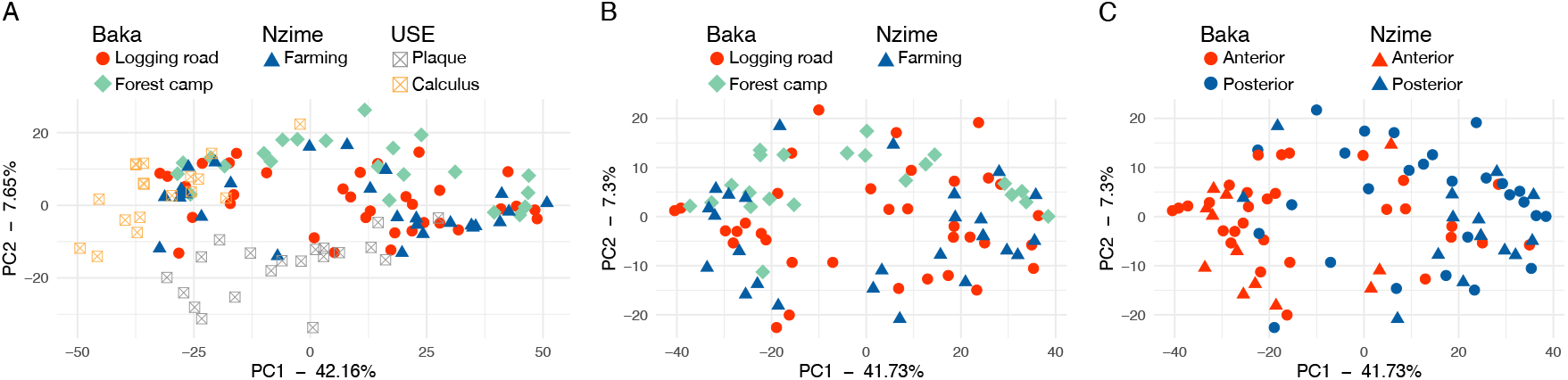
Species community structure of dental plaque from Baka and Nzime differs by tooth sampling location and oxygen tolerance. **A**. PCA of microbial communities from Baka, Nzime, industrial dental plaque and industrial calculus show the structure of these groups is different. Points colored and shaped by village location, which corresponds to ethnic group, or industrial sample type. **B**. PCA of Baka and Nzime samples with points colored and shaped by village location, which corresponds to ethnic group. **C**. Same as (B) with shapes corresponding to ethnic group and colors corresponding to tooth collection site.

When considering variation only within BN dental plaque, the samples appear to form two clusters along PC1, with no distinction by village, market integration, ethnic background, age group, or sex (Figure 6B, Supplemental Figure S13). Instead, they separate by tooth site (anterior *vs*. posterior teeth, Figure 6C), which is significant by PERMANOVA (F =13.0, R^2^ = 0.141, p = 0.001). This suggests that the oral microbiome profile of Baka and Nzime are more strongly influenced by differences in oral environment than by ethnic group, lifestyle factors, or diet.

To determine the species most influencing compositional differences between sample types, we performed differential abundance analysis focused on the differences between USE and BN samples, as well as between anterior and posterior samples of BN dental plaque. Compared to USE dental plaque, the BN dental plaque samples were enriched in anaerobic species, while compared to USE calculus, the BN dental plaque samples were enriched in aerotolerant species (Figures 7A,B, Supplemental tables S7, S8). This indicates that the biofilm maturation stage of BN dental plaque is intermediate between that of USE dental plaque and USE calculus, which may be related to the absence of frequent biofilm reduction through toothbrushing. Within Baka and Nzime dental plaque, anterior samples were enriched in aerobic and facultatively anaerobic taxa compared to posterior samples (Figure 7C, Supplemental table S9), which may be driving the separation of these sample types in the PCA. From this distinction, there appears to be a gradient of species oxygen tolerance that aligns with the sampling site, where anterior samples have a profile characterized by more aerotolerant species, and posterior samples have a profile characterized by more anaerobic species. This is consistent with anterior teeth having greater exposure to air during breathing, speaking, and eating compared to posterior teeth.

**Figure 7.**
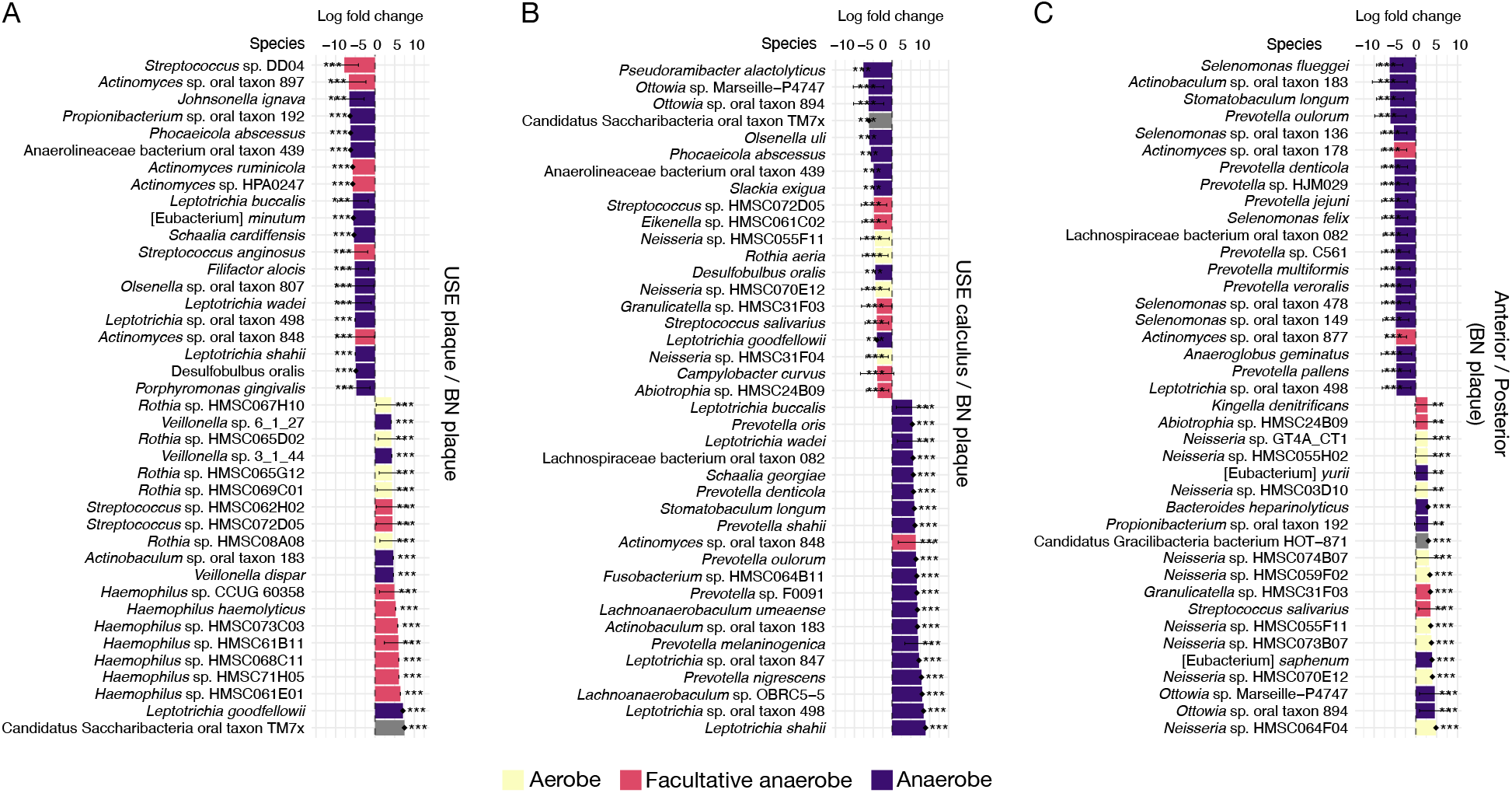
Differentially abundant taxa suggest different levels of oxygen tension in USE and BN dental plaque biofilms. Bars are colored by aerotolerance of each species. **A**. Top 40 species enriched in USE dental plaque (positive values) and BN dental plaque (negative values). **B**. Top 40 species enriched in USE calculus (positive values) and BN dental plaque (negative values). **C**. Top 40 species enriched in BN dental plaque anterior collection sites (positive values) and posterior collection sites (negative values). * p < 0.05, ** p < 0.01, *** p < 0.001; dots on bars indicate structural zeros determined by ANCOM-BC.

### Functional characteristics of Baka and Nzime dental plaque

While the taxonomic composition of dental biofilms among the Baka, Nzime, and USE populations were similar, oral taxa may have large pangenomes ^50–52^, and it is possible that there are distinctive differences in the gene content of the taxa present. We used HUMAnN3 functional classification to generate a metabolic pathway abundance table for functional analysis. The amount of data that could not be classified by HUMAnN3 was significantly higher in the BN samples compared to USE dental plaque, suggesting more uncharacterized and uncategorized gene content in the BN dental plaque samples (Supplemental figure S15A). This is reflected in the lower number of pathways detected in BN plaque compared to USE dental plaque and USE calculus (Supplemental figure S15B, C). To determine if the metabolic pathways distinguish sample groups more distinctly than taxonomic composition, we performed a PCA on the metabolic pathways (Figure 8). Sample group separation based on metabolic pathway content was less distinct between BN dental plaque, USE dental plaque, and USE calculus (Figure 8A) than by species profiles (Figure 6). However the BN dental plaque again separates along PC1 by anterior/posterior tooth collection site, which is significant by PERMANOVA (R^2^ = 0.114, F = 10.5, p = 0.001), rather than by ethnic group or village (Figure 8B,C, Supplemental figure S15), similar to the species profile. Repeating the analysis with the gene families produced a similar sample clustering (Supplemental figure S16).

**Figure 8.**
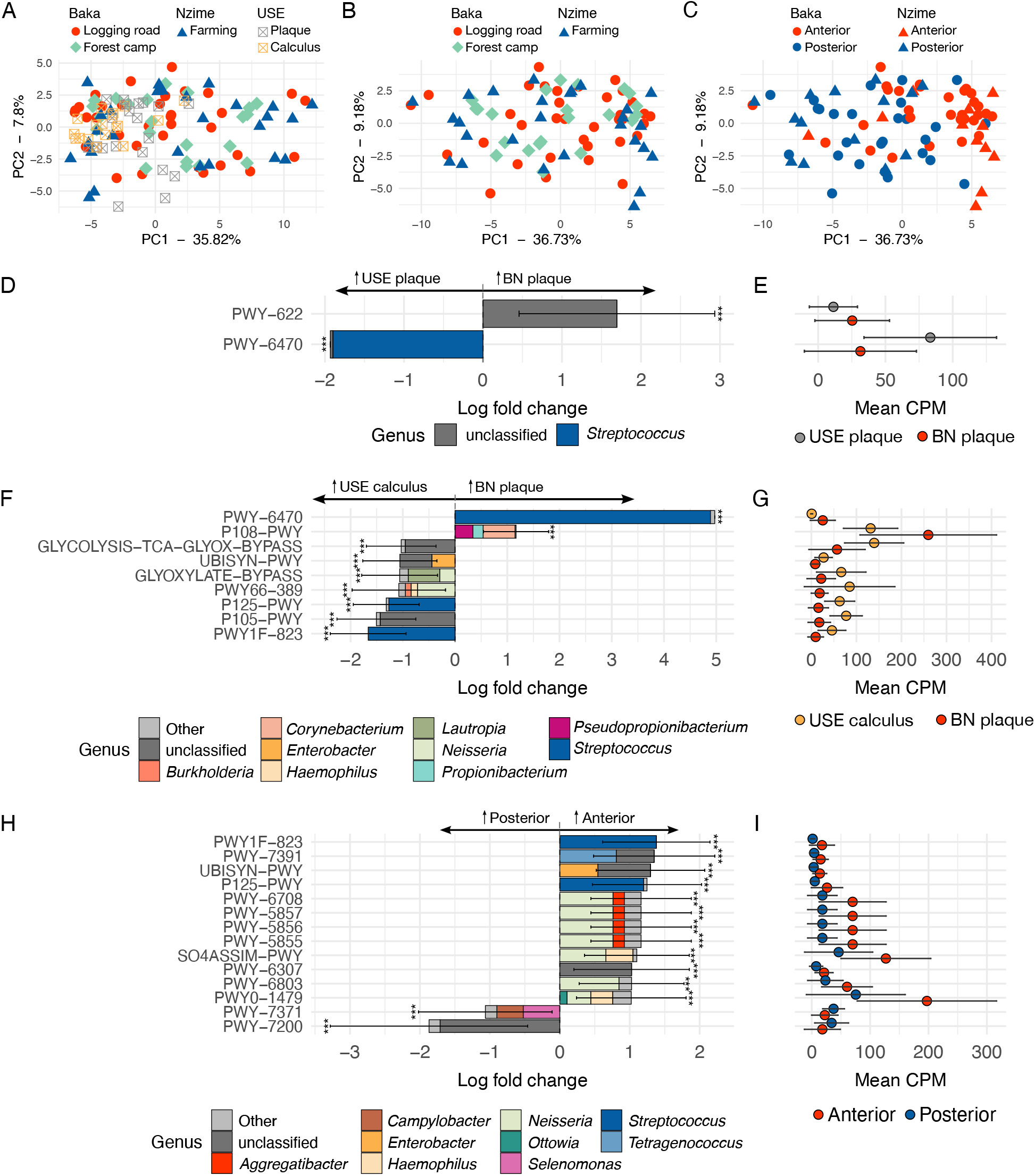
Functional differences of dental plaque communities from Baka, Nzime and industrial dental biofilms are not clearly delineated. **A**. PCA of metabolic pathway content from Baka, Nzime, USE dental plaque, and USE calculus show the pathway content is similar between these oral microbiome sources. Points colored and shaped by market integration, corresponding to ethnic group, and industrial sample type. **B, C**. PCA of metabolic pathway content in BN samples with points colored and shaped by market integration (**B**) and tooth collection location (**C**). **D, F, H**. Metabolic pathways that are 1-fold more or less abundant between groups are shown. Colors indicate the proportion of the metabolic pathway attributed to each genus present at ≥ 5% abundance in each sample group. **E, G, I**. Mean copies per million (CPM) per sample group of the differentially abundant metabolic pathways. **D, E**. BN samples compared to USE dental plaque samples. **F, G**. BN samples compared to industrial calculus samples. **H, I**. Anterior samples compared to posterior samples of Baka and Nzime individuals. *** q < 0.001.

We assessed whether there are distinct metabolic pathways that characterize the sample groups by performing differential abundance analysis. No clear patterns were discernible in the types of metabolic pathways that characterized each group (Figures 8D-I), with one exception. Within BN dental plaque, anterior tooth samples were enriched in pathways related to oxidation-reduction reactions (Figure 8H,I) including PWY-5855, PWY-5856, PWY-5857,PWY -6708, PWY-7371, UBISYN-PWY, and SO4ASSIM-PWY (Supplemental tables S10-S12), which may be involved in maintaining a reduced environment for the biofilm in a location with higher oxygen exposure.

### Identification of additional taxonomic diversity in Baka and Nzime dental plaque

Previous studies suggest that there is diversity in human oral and intestinal microbiomes that is not currently represented in databases that are typically used for taxonomic profiling, such as those maintained by the NCBI ^53,54^. This trend is also indicated in the BN dental plaque samples analyzed here by the lower percentage of reads assigned taxonomy (Supplemental figure S7) and the lower number of classified metabolic pathways and genes (Supplemental figures S15, S16). To determine whether the dental plaque of Baka and Nzime individuals contain uncharacterized taxonomic diversity, we performed an additional taxonomic profiling using a database containing metagenome-assembled genomes (MAGs) that include taxa not represented in NCBI RefSeq from Passoli, *et al*. ^53^. We found that more reads were assigned taxonomy when using a database that includes MAGs than when using a database restricted to RefSeq genomes alone (Figure 9A), although the increase within each group was small and not significant.

**Figure 9.**
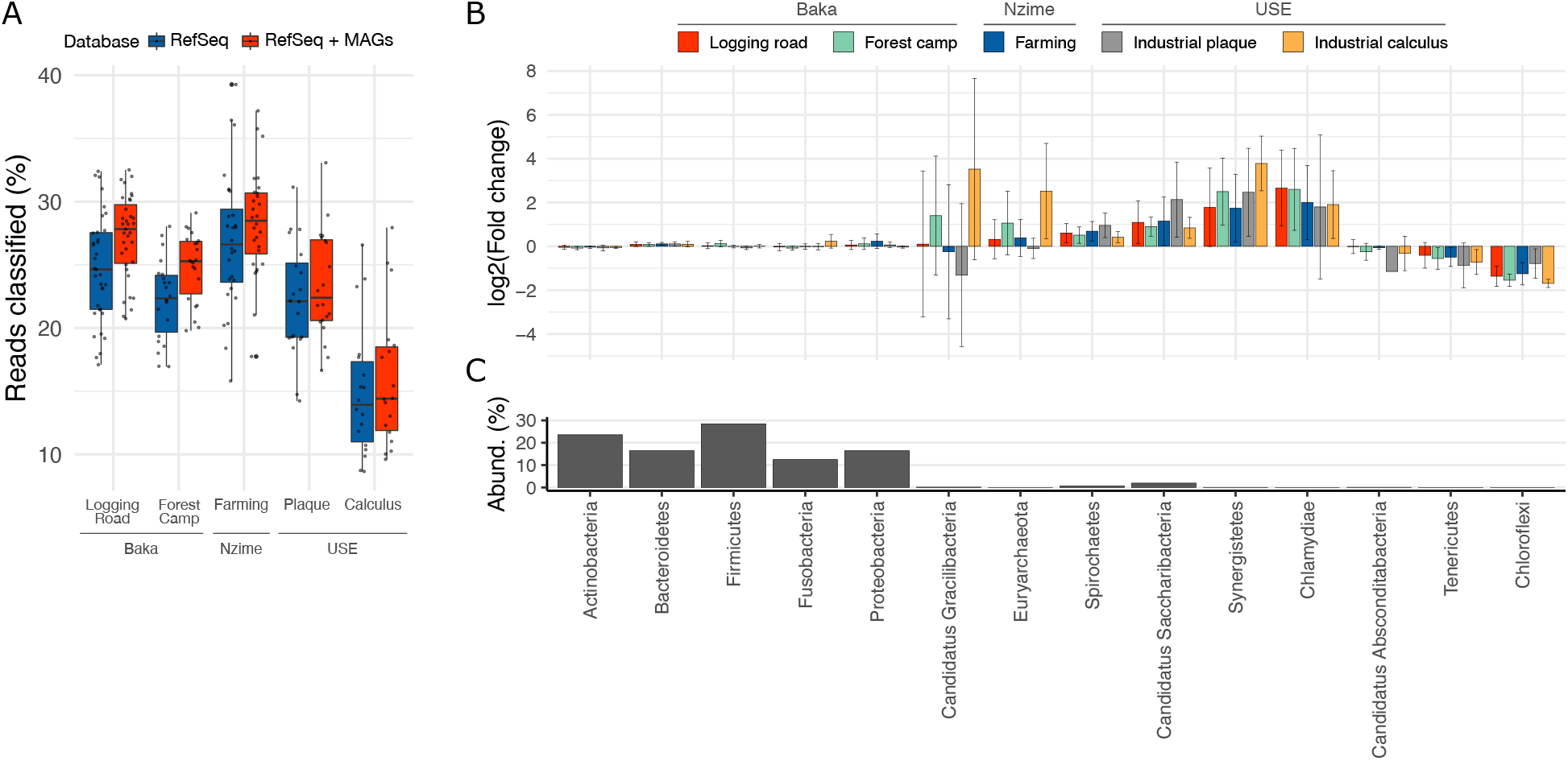
Novel taxa are present in dental plaque samples regardless of market integration status. **A**. The percent of reads assigned taxonomy increases per sample when using a database that excludes (RefSeq) or includes (RefSeq + MAGs) metagenome-assembled genomes (MAGs). **B**. Enrichment or depletion of taxa in the 14 phyla known to exist in the mouth, when profiled with a database excluding MAGs vs. including MAGs. **C**. Relative abundance in supragingival dental plaque of the 14 phyla known to exist in the mouth, averaged from 3 studies whose abundance-based tables are cataloged on the Human Oral Microbiome Database (homd.org). The 5 phyla with least change in abundance between databases (**B**) are the most abundant (**C**).

To determine if there were substantial changes in the proportions of known oral taxa identified by profiling with a database including MAGs, we looked at the difference in proportion of assignments to the 14 phyla previously described in the oral cavity: Actinobacteria, Bacteroidetes, Candidatus Gracilibacteria (formerly GN02), Candidatus Absconditabacteria (formerly SR1), Candidatus Saccharibacteria (formerly TM7), Chlamydiae, Chloroflexi, Euryarchaeota, Firmicutes, Fusobacteria, Proteobacteria, Spirochaetes, Synergistetes, and Tenericutes ^5^ (Figure 9B). Five of the phyla, Actinobacteria, Bacteroidetes, Firmicutes, Fusobacteria and Proteobacteria, which include the most prevalent and best-characterized oral taxa, showed little difference in their abundance between profiling with the two databases for all sample groups. These are also the most abundant phyla in supragingival dental plaque (Figure 9C). Candidatus Gracilibacteria and Euyarchaeota showed high variability across samples in each group. Four of the phyla that include less-well characterized oral taxa were uniformly more abundant when profiling with a database including MAGs (Spirochaetes, Candidatus Saccharibacteria, Synergistetes, and Chlamydiae), while Candidatus Absconditabacteria,Tenericutes and Chloroflexi were uniformly less abundant when profiling with a database including MAGs. These results indicate that there is a similar set of taxa being missed in both USE and BN samples, and that these are largely present in the 9 less-abundant and less-well studied phyla occupying the oral cavity.

## Discussion

The dental plaque microbiome shows high conservation across populations with distinct diets, lifestyles, and different levels of market integration. While the range of variation in species profiles of BN dental plaque samples does not fall completely within that of previously described USE oral biofilms (dental plaque and dental calculus), the dominant taxa are largely the same as those in North American and European populations. This is in contrast to the gut microbiome, which has been shown to have a highly variable and distinct microbial community structure between groups that consume industrially-processed diets and those that rely more heavily on hunting, foraging, or small-scale farming ^6,8–10,14,55^. Our results highlight that the distinct characteristics of microbiomes from independent body sites cannot be broadly generalized as pertaining to “the human microbiome”.

While several studies have reported pronounced differences between stool microbiomes of North Americans and Europeans compared to those of individuals with lower market integration from southwest Cameroon and neighboring Central African Republic, previous studies of saliva found differences that were less pronounced ^9,10,14^. We found that in dental plaque differences are even less apparent. Instead, dental plaque microbial composition is primarily structured by oxygen tolerance. The high degree of conservation between BN dental plaque, and between BN dental plaque compared to USE dental biofilms, supports that diet has only a limited influence on the dental plaque microbiome. As the major nutrient source for many oral microbes is salivary mucins ^56^, diet-based fluctuations in oral species abundance may be a minor contributor to overall species profiles. In this way, the oral cavity contrasts sharply with the gut, where digestive passage times are considerably longer and host diet is the major nutrient source for taxa that make up the stool microbiome. Although microbes colonizing gut mucosa, in contrast to stool, have not been extensively studied in diverse populations, we speculate that they may likewise show reduced variation across populations with distinct diets and lifestyles due to their higher reliance on host mucins as a primary source of nutrition ^57^.

Dental plaque microbiomes are notably different between healthy teeth and those experiencing dental disease, particularly dental caries or periodontal disease ^58,59^. Therefore, it is important for studies of oral microbiomes to take into account study participant dental health. We found similar rates of dental pathologies between Baka and Nzime individuals, so it is unlikely that there are microbial composition differences between the two groups stemming from differences in dental health. However, there are clinical aspects that could be investigated in future studies, such as site-specific microbial profile differences, or subgingival vs. supragingival plaque profiles, which this study was not designed to investigate. These may provide insight into processes of dental biofilm development that sustain health or promote disease, particularly in the absence of regular oral hygiene.

The overall species profile of BN dental plaque samples appears to be intermediate between USE calculus and USE dental plaque, a point of caution for future comparative studies. While the most abundant dental plaque taxa of the Baka and Nzime individuals are all well-known oral taxa, they are enriched in anaerobic taxa compared to industrial dental plaque but enriched in aerobic taxa compared to industrial calculus. The distinction in aerotolerance of differentially abundant taxa suggests the BN dental plaque biofilms are in a more mature stage of biofilm development than is typical in dental plaque from groups with heavy market integration. This may be due to differences in daily oral hygiene regimens that reduce dental biofilm buildup (i.e., toothbrushing). As dental plaque biofilms develop and mature according to a predictable species succession ^60^, the maturation stage can be distinguished by species composition. Thus, hygiene practices and biofilm maturation stage are important factors that need to be considered in future dental plaque studies.

It has been proposed that some microbial taxa present in populations with limited market integration may preserve diversity that has been lost in heavily integrated populations ^6,61^. We present evidence that the dental biofilm communities of not only Baka and Nzime individuals, but also North American and European individuals, contain taxonomic diversity not currently represented in existing databases derived from NCBI. This suggests that the mouth, in addition to the gut, may be able to sustain higher levels of diversity than can currently be identified; however, the unidentified taxa may have similar prevalence across a range of market integration levels.

It will be of interest in future studies to identify the taxa in oral samples that are not presently in our databases, to characterize their genomic content, and to determine if they are associated with oral health or disease. Further, description and characterization of novel taxa, especially at the strain-level, may help us study co-evolution of microbial species and communities through deep time using ancient DNA (aDNA) ^62^. One major question faced by ancient metagenomics researchers is whether the microbial taxa currently present in databases are sufficient representatives of those in ancient samples. Including taxa from populations with low levels of market integration in databases for profiling ancient microbiomes may improve taxonomic representation, and therefore taxonomic identification, and aid in clarifying the extent of historical microbiome diversity.

Here we show that subsistence strategy minimally shapes the dental plaque microbiome. While there are small differences in species profiles between dental plaque from groups with high and low levels of market integration, they appear to be largely related to biofilm maturation and dental hygiene practices that delay maturation (i.e., toothbrushing). Consequently, dental calculus, which represents a more mature and less disturbed oral biofilm, may be a more appropriate comparative material for future studies of dental biofilms from populations with low market integration. We show that diet and lifestyle factors influence dental plaque microbial communities to a much lower extent than they do the gut microbiome, indicating that different forces shape the taxonomic composition at these two body sites. Such distinctions have implications for understanding how human microbiomes respond to industrialization-related changes (diet, hygiene, urbanization) and how this relates to health and disease - both in populations currently undergoing these changes, and in populations that experienced them in the past.

## Materials and Methods

### Samples and study populations

This study was conducted as part of a broader investigation of foraging strategies among the Baka people of southeastern Cameroon, with a specific focus on the importance of plant dietary resources in forested tropical environments ^38,63^. For details see Supplemental Methods. Dental plaque samples and metadata were collected by author S.G. between October-November 2018, with the assistance of a Nzime research assistant and interpreter who spoke the local dialects of both studied populations. This occurred during the major rainy season, when many Baka individuals reside in forest camps and participate in major hunting and wild edible-foraging excursions in the forest ^38^. A total of 47 individuals (30 Baka and 17 Nzime) from five different communities (three Baka and two Nzime villages) in the Messok and Lomie districts of the Haut Nyong Division of southeastern Cameroon participated in the study. These Baka communities and Nzime villages were located along the logging road, with the exception of one Baka forest camp, which was located at a distance of about an hour and a half by foot from the logging road. Based on location, access to the market economy and to health and school services differed among the communities in this study.

### Informed consent and ethical review

Before the onset of the research we received the Free Prior Informed Consent from both the local authorities of the villages and the participants. The study received the research permit from the Ministry of Scientific Research and Innovation (MINRESI n°000115/MINRESI/B00/C00/C10/C12 and n°000116/MINRESI/B00/C00/C10/C12). Members of Baka and Nzime communities were contacted by project members about the possibility of participating in a study on dental health and diet. Prior to sample and metadata collection, the study and its goals and methods were explained to prospective participants in the five communities. Study volunteers provided informed consent and agreed to a dental health assessment and dental cleaning, followed by an analysis of their dental biofilm. To ensure representation of the general adult population as well as age and gender balance in the study, we considered all community adults to be eligible, including individuals with missing teeth, periodontitis, and visible caries. Participant demographic metadata and oral health status (as determined at the time of the dental cleaning) were recorded for each individual (Supplemental figure S1). Individuals who had taken antibiotics within the past 6 months were excluded from the study. None of the women who participated were pregnant. After the completion of the study, project member S.G. met with local authorities and village members in different meetings to share the results of the study and distribute a booklet describing a summary of the findings. The study design was approved by the Ethical Committee from the Ministry of Health of Cameroon (n°2018/06/1049/CE/CNERSH/SP) and the ethics committee of Leipzig University (196-16/ek). It also adheres to the Code of Ethics of the International Society of Ethnobiology ^39^.

### Sample collection and storage

Sample collection proceeded as follows: first, the participants’ teeth were brushed with a dampened toothbrush to remove food debris and loosely attached dental plaque. Next, the overall condition of the teeth was visibly inspected, and decayed, missing, filled or altered teeth were recorded, along with any instances of loose teeth, bleeding gums, and teeth with visible root exposure (Supplemental Table S1, using the FDI World Dental Federation notation). Using a sterilized (autoclaved) dental scaler, we collected two biofilm samples from each individual, one from the anterior teeth (incisors or canines, depending on availability) and one from the posterior teeth (premolars and molars, again where available). Each biofilm sample was placed onto a piece of sterile gauze, dried, and individually packaged into small sealed plastic bags containing a silica gel desiccant to reduce humidity. The dried biofilm samples were stored at ambient temperature (10-28°C) for several weeks until the completion of fieldwork, after which they were transported to the laboratory facility at the Max Planck Institute for the Science of Human History (now the Max Planck Institute for Evolutionary Anthropology), where they were stored at −20°C.

### DNA Extraction and Next Generation Sequencing

DNA was extracted from samples using the Qiagen PowerSoil kit. For each sample, a small piece of gauze with visible dental plaque was cut from the main piece and placed in a PowerSoil bead tube. Two controls were included for each extraction batch: a gauze control and an empty tube control. Both controls were processed alongside the samples throughout the entire extraction and library building process. For details see the Supplemental Methods. DNA was extracted following the detailed protocol “DNA Extraction from Modern Dental Plaque on Gauze” ^64^.

Illumina library preparation followed the detailed protocol “Non-UDG treated double stranded ancient DNA library preparation for Illumina sequencing” ^65^, and included a library blank control, which used water as input instead of sample. The libraries were indexed following the detailed protocol “Illumina double-stranded DNA dual indexing for ancient DNA” ^66^, adapted from ^67^, with the modification that the entire process was performed in a modern DNA lab. All libraries were sequenced on an Illumina NextSeq500 with 2×150bp chemistry. Each library was sequenced on 2 or 3 flow cells to a depth of approximately 10 million reads per sample (Supplemental Table S1). Sequencing data was demultiplexed with an in-house pipeline.

### Comparative Data

Previous oral microbiome studies of non-industrial populations have focused primarily on saliva, and to a lesser degree buccal mucosa. Saliva, however, is not a true microbiome but rather is formed through the shedding of microbial cells from multiple distinct oral sites ^1,2^. To our knowledge, no studies of dental plaque from non-industrial populations have been published to date. For this study, we selected as comparative datasets dental plaque and dental calculus from industrialized populations, buccal mucosa samples from both non-industrialized and industrialized populations, and saliva from non-industrialized and industrialized populations. For industrial food consumers, both dental plaque and dental calculus were included because they have been shown to contain distinct species profiles ^41^. The six comparative datasets consisted of: (1) 20 dental plaque samples of industrial food consumers in the United States from the Human Microbiome Project (HMP) (PRJNA48479) ^1^; (2) 18 dental calculus samples of industrial food consumers in Spain (PRJEB31185 and PRJEB34569) ^30,41^; (3) 14 buccal mucosa samples of Yanomami hunter-gatherers in the Amazon rainforest of Venezuela (PRJNA245336) ^18^; (4) 14 buccal mucosa samples from industrial food consumers in the United States from the HMP (PRJNA4847) ^1^; (5) 24 saliva samples from hunter-gatherers and traditional farmers in the Philippines (PRJEB14383) ^20^; and (6) 5 saliva samples from industrial food consumers in the United States from the HMP (PRJNA4847) ^1^. A full list of sample accession IDs is provided in Supplemental Table S1. The Yanomami comparative dataset (Clemente, et al. 2015) contained the only samples that were treated with multiple displacement amplification (MDA) after extraction. Directly comparing the microbial species profiles of dental plaque, buccal mucosa, and saliva of industrial and non-industrial populations allowed us to investigate whether stronger differences exist between oral sites or between industrial and non-industrial populations.

### Data Processing

All sequencing data produced for this study, as well as the comparative datasets, were processed with the nf-core/eager pipeline ^68^. This included adapter trimming, read merging, and quality filtering with AdapterRemoval ^69^, and mapping against the human genome (HG19) with bwa aln -n 0.02 -l 1024 ^70^ to remove human reads prior to analysis. All sequencing files for each library were concatenated during the eager run to create a single fastq file per library, which was mapped against the human genome. All reads that did not map to the human genome were used for taxonomic profiling.

### Taxonomic Profiling and Clean-up

For taxonomic profile exploration, we used MALT v.0.4 ^71^ within the nf-core/eager pipeline, using a RefSeq database (RefSeq_malt) that included bacteria, archaea, and the human genome ^30^. To assess post-collection environmental contamination, sample integrity was checked with the R package cuperdec (https://github.com/jfy133/cuperdec) ^30^. To remove possible contaminants introduced to the samples during laboratory processing, we used the R package decontam ^72^. All species that were identified as contaminants were removed from the species table before performing further analyses. We tested for batch effects in our extraction data by performing diversity analyses and comparing the extraction batches.

### Alpha-diversity

To assess within-sample diversity, we calculated the number of observed species and the Shannon index. Taxa present at less than 0.001% abundance were removed from the table, as recommended for MALT ^73^, which produces a high number of low-abundance false-positive taxonomic assignments. Heatmaps of the top 50 most abundant species were generated by determining the 50 most abundant species in each group (Baka and Nzime dental plaque, industrial dental plaque, and industrial dental calculus) and then these species were subsetted from the species table.

### Factor Analysis

Exploratory factor analysis was performed to reduce the number of variables (species) for beta diversity analyses, rather than using an abundance-based cut-off, with the R package psych ^74^ with oblimin rotation and minres factoring method. A species table containing only the species that were in the first factor, minimum residual 1 (MR1), which explains a majority of the variation in the data, was used for further diversity analyses. Heatmaps of the CLR-transformed read counts for each species in MR1 were generated with ggplot (Figures S11, S12).

### Beta-diversity

Beta-diversity analyses were performed on a species table including only the species in minimum residual 1 from the factor analysis. A value of +1 was added to all species read counts, which were then CLR-transformed and a principal components analysis (PCA) performed using the R package mixOmics ^75^, and plots generated with ggplot. To determine if sample groups in the have significantly different species profiles, a PERMANOVA was performed with the function adonis2 from the R package vegan ^76^. We additionally separated the samples by anterior and posterior tooth collection site, as this difference was the strongest explanation of variation between groups in the PERMANOVA, and repeated the beta-diversity analyses on only anterior and only posterior samples.

### Differential abundance

Species differential abundance was tested with ANCOM-BC ^77^. ANCOM-BC was performed following the examples provided on the ANCOM-BC github page, and plots were generated with ggplot.

### Functional Profiling with HUMAnN3

We performed functional profiling of all samples with HUMAnN3 ^78^ with default settings. We used the path abundance output table with paths classified with MetaCyc pathways ^79,80^, and the gene families output table classified with UniProt90, both normalized by copies per million (CPM). For pathway and gene family analysis we selected the total abundance lines from the eache table (this removed all species-level assignments) for diversity analyses. The variation in pathways and gene families between groups were assessed using PCA and differential abundance following the same steps as for species analysis, see Supplemental Methods for details.

### Taxonomic profiles with and without metagenome-assembled metagenomes

To determine if there are taxa present in the Baka and Nzime samples that are not represented in the NCBI RefSeq database, we used Kraken2 ^81^ and built two databases, one containing only RefSeq genomes (here, RefSeq_kraken) and the other containing those same RefSeq genomes and the metagenome-assembled genomes published by Pasolli et al. ^53^ (here, RefSeq_Pasolli_kraken). Taxonomic profiling using Kraken2 was applied to all Baka and Nzime dental plaque, industrial dental plaque and industrial calculus datasets for both databases. The number of reads that were classified and unclassified in each sample in each run were determined from the Kraken2 output file, and a Wilcox rank sum test was used to determine if there was a significant difference between the databases for the number of assigned reads in each group. To determine if there are differences in the number of reads assigned to known oral taxa between the two databases, we calculated the log fold change in relative abundance of the 14 phyla described in the human oral cavity ^5^ when profiling with RefSeq_Pasolli_kraken vs RefSeq_kraken. For details see Supplemental Methods.

### Taxonomic ratios

We calculated the ratio of the relative abundance of the phyla Bacteroidetes vs. Firmicutes and the ratio of the relative abundance of the genera *Prevotella* vs. *Bacteroides* within the Baka and Nzime samples, as well as within industrial dental plaque and calculus from the MALT-generated species profiles.

## Acknowledgements

We thank Ernest Simpoh for assistance with interpretation during participant recruitment and sample donation, Antje Wissgot for the sequencing assistance of the dental plaque samples, and Alexander Hübner for constructing the Kraken2 databases. This research was supported by the European Research Council under the European Union’s Horizon 2020 research and innovation program, grant agreement number STG–677576 (“HARVEST”), the Deutsche Forschungsgemeinschaft (DFG, German Research Foundation) under Germany’s Excellence Strategy (EXC 2051, Project-ID 390713860), the Werner Siemens Stiftung (“Paleobiotechnology”), and the Max Planck Society.

## Data availability

All raw sequencing data is deposited in ENA under the project code PRJEB42572. All scripts for analysis can be found in the github page for the project https://github.com/ivelsko/Cameroon_plaque.

## Supplemental Information

## Supplemental Methods

### Sample Populations

Southeastern Cameroon has a tropical humid climate, and Baka and neighboring Nzime villages are situated within evergreen and moist semi-deciduous forests at an elevation of 300-600 meters and with an average temperature of 25°C. There are two rainy and dry seasons per year, and annual precipitation can reach 1500 mm ^1^. Baka and Nzime villages are distant from urban-industrialized centers, being located approximately eight hours away by car from the capital of Yaoundé, of which four hours require travel on unpaved logging roads.

In this study, we partnered with members of two distinct ethnic groups, the Baka, a Ubangian speaking people who practice foraging and small-scale cultivation, and the Nzime, a Bantu-speaking people who practice subsistence-level agriculture ^2,3^, to better understand the influence of diet and lifestyle on oral microbial communities. Although living in the same area, Baka and Nzime lifeways greatly differ. The Nzime rely on subsistence agriculture, cultivating mostly cassava and plantains as major crops, and also producing cacao for trade. The Baka rely on a more mixed subsistence strategy, combining foraging (hunting, gathering, and fishing) with small-scale farming ^4^. The Baka and Nzime have long had an economic relationship based on the exchange of foraged forest and farmed agricultural products, but over the past 60 years the Baka have become more sedentary and reliant on local agricultural products, and they increasingly work as agricultural laborers in Nzime fields.

The diets of both the Baka and the Nzime are primarily plant-based, with a focus on cassava and plantains, supplemented by bushmeat and wild or cultivated vegetables. Both the Baka and Nzime have limited access to market economy goods through small local shops and traders traveling along logging roads, and such goods consist mostly of machetes, metal cookpots, salt, rice, sugar, flour, sardines, tomato salsa, bouillon cubes, candies, and alcohol. Toothpaste is not typically available, and dental cleaning is uncommon, particularly among the Baka. While the main components of Baka and Nzime diets overlap, Baka diets have a higher degree of dietary seasonality and they consume a wider diversity of wild products in their daily meals, although such diversity tends to decrease with higher integration into the market economy ^4,5^. The Nzime keep domestic animals, while the Baka do not. Due to their higher income from agricultural surplus, the Nzime have greater access to market goods and integrate a higher proportion of processed foods obtained from local shops into their diet.

### Dental Health Assessment Overview

Samples were scored for presence/absence of associated pathologies based on entries in the decayed, missing, root visible, and altered fields (Supplemental table 1). No members of the Baka or Nzime communities had dental fillings (or other hardware such as crowns or dentures). All noted health conditions on anterior teeth, both in the maxilla (tooth numbers 11, 12, 13, 21, 22, 23) and mandible (tooth numbers 31, 32, 33, 41, 42, 43), were recorded collectively for the anterior dental plaque sample, and all health conditions on posterior teeth, both in the maxilla (tooth numbers 14-18, 24-28) and mandible (tooth numbers 34-38, 44-48), were recorded collectively for the posterior dental plaque sample. Dental health summary plots are provided in Supplemental Figures S2 and S3 and in the R markdown file on the github page 02-scripts.backup/Cameroon_taxonomy_oral_health.Rmd.

Samples were scored for presence/absence of associated pathologies based on entries in the Decayed, Missing, Root_visible, and Altered fields. that had at least 1 tooth entered were converted to “Yes” and all cells with NA were converted to “No”, to get a binary distinction between dental plaque collected from an area with any oral pathology and dental plaque collected from an area with no pathology. The number of samples with pathology and without pathology (both anterior and posterior), “Yes” and “No” respectively, were counted for different metadata categories, and the counts plotted. No statistical testing was done to determine if there are statistically significant differences because this study was not designed to detect statistical significance, and is descriptive instead.

### DNA Extraction Blanks

DNA was extracted from samples using the Qiagen PowerSoil kit in 7 in batches, and each batch was processed with a gauze control and a blank tube control. For the gauze control, one sample was randomly selected from the batch, and a piece of gauze with no visible dental plaque, far from the sampled dental plaque location, was cut and placed in a PowerSoil bead tube. The gauze control was used to assess the potential presence of non-oral environmental microbes on the sample collection material, as this can bias oral sample profiles ^6^. However, because the gauze control was obtained from a larger piece of gauze also containing a plaque sample, it is possible that trace oral bacteria might be later detectable on some gauze controls, and this was assessed during data analysis. The empty tube control was used to assess potential laboratory background contamination and consisted of a PowerSoil bead tube that contained no gauze. From one Baka individual, both the anterior and the posterior plaque produced too little DNA to be successfully built into libraries, so 45 of 46 individuals sampled were included in the study.

### Taxonomic Profiling and Clean-up

To create a species table for analysis, the .rma6 files produced by MALT were uploaded to MEGAN6 ^7^ using the compare function, and tsv files for summed species-level read count assignments were exported. The total proportion of reads assigned taxonomy in each sample ranged from 50%-90% (Supplemental Figure S7). For preservation assessment, we used the R package cuperdec (https://github.com/jfy133/cuperdec) ^8^ to determine the unweighted proportion of taxa in each sample that have an oral source. Although designed to assess preservation of ancient dental calculus samples, it can also indicate if modern samples are well-preserved, or if they have been overgrown by environmental contamination. We used the adaptive burn-in filter with a percent threshold cut-off of 53. All steps for cuperdec analysis can be found in the R markdown file on the github page 02- scripts.backup/Cameroon_taxonomy_cuperdec.Rmd, which uses tidyverse formatting ^9^.

To identify taxa that may derive from laboratory contamination, we used the R package decontam ^10^. This package uses the profiles of samples and blanks to identify species that are likely to originate from contamination. Prior to running decontam, we identified samples that resembled controls, and controls that resembled samples, by performing a PCA using the R package mixOmics after adding +1 to each value and performing a centered-log ratio (CLR) transform, and plotting the results. Four of the six gauze controls plotted closely to the samples, and one sample plotted with the blanks (Supplemental figure S10). The four gauze controls contained trace oral bacterial contamination, likely saliva from sample collection, while the sample plotting with blanks had insufficient biological material for analysis and very low read counts. These four gauze controls and one sample were removed from the species table before running decontam in order to avoid skewing the contamination profile for decontam analysis, and they were also removed from all further analyses. Within decontam, we selected the combined frequency and prevalence method and a threshold of 0.9 for contaminant identification with decontam. All steps for decontam analysis can be foundin the R markdown file on the github page 02-scripts.backup/Cameroon_taxonomy_decontam.Rmd, which uses tidyverse formatting. We additionally checked the samples for batch effects, and found no significant differences between batches (Supplemental Figure S6). All steps for batch effect analysis can be found in the R markdown file on the github page 02-scripts.backup/Cameroon_taxonomy_batches.Rmd, which uses tidyverse formatting.

### Alpha-diversity and species abundance heatmaps

Statistical testing of differences between groups for alpha-diversity metrics was performed using the package rstatix ^11^. Raincloud plots were generated following the tutorial by Cedric Sherer (https://www.cedricscherer.com/2021/06/06/visualizing-distributions-with-raincloud-plots-and-how-to-create-them-with-ggplot2/). All steps for alpha-diversity analysis can be found in the R markdown file on the github page 02-scripts.backup/Cameroon_taxonomy_alphadiv.Rmd. Alpha diversity analyses were performed in R using the packages vegan ^12^, compositions ^13^, and janitor ^14^.

For plotting the heat map of abundant species, the read counts for each were adjusted by adding +1 followed by CLR-transformation. The species were hierarchically clustered using the function hclust from the R package stats ^15^ and then plotted in a heatmap with ggplot.

### Factor Analysis

MALT identifies a high number of false-positive low-abundance species ^16^, which need to be removed prior to analyses. Rather than using an abundance-level cut-off to remove these false-positive species, we used factor analysis to select the species that most strongly distinguish sample groups, meaning those with highest loadings in the first factor, which explains the largest proportion of variation in the dataset, and worked instead with only these species. Factor analysis was performed individually for a table with BN plaque, USE plaque, and USE calculus samples, and identified 123 species in the first minimum residual (factor). Factor analysis was performed again for a table of only Baka and Nzime samples, and identified 117 species in the first minimum residual (factor). A discriminant analysis was performed with the R package MASS ^17^ to test the ability of the species in the first factor to discriminate various metadata groups. All steps for factor analysis can be found in the R markdown file on the github page 02- scripts.backup/Cameroon_taxonomy_discrimspecies.Rmd.

### Beta-diversity

Beta-diversity analysis was performed using principal components analysis (PCA). The number of factors to include in the PCA was determined by visual examination of the scree plot, where all factors that preceded a large drop in percent variation explained were kept. In most cases this was 2 or 3 factors. To determine which species were contributing to the separation of groups in the plot, a bi-plot containing the 10 species with the strongest loadings in PC1 positive and negative values was generated (20 species in total). All steps for beta diversity analyses can be found in the R markdown files on the github page 02- scripts.backup/Cameroon_taxonomy_betadiv.Rmd. All steps for beta diversity analyses on samples separated by anterior/posterior tooth collection site can be found in the R markdown files on the github page 02-scripts.backup/Cameroon_taxonomy_anterior_div.Rmd, 02- scripts.backup/Cameroon_taxonomy_posterior_div.Rmd.

### Differential abundance

All steps for ANCOM-BC differential abundance analysis can be found in the R markdown files on the github page 02-scripts.backup/Cameroon_taxonomy_ancom-bc_DA_species.Rmd.

### Functional Profiling with HUMAnN3

The pathway table was subjected to decontam to remove entries that were identified because they were introduced by laboratory contaminants. A factor analysis and linear discriminant analysis were performed on the pathway table, using the same steps as for the species table. To perform PCA, a table with only the pathways in the first minimum residual was used, and a value of +1 was added to all CPM, the table was CLR-transformed, and PCA was performed with mixOmics and plotted with ggplot. Significant differences between groups were assessed by PERMANOVA with the function adonis2 from the package vegan. Plots were generated in ggplot.

We additionally determined differentially abundant pathways using ANCOM-BC ^18^. To consider a pathway differentially abundant, we required that the log fold change between groups be ≥1, with a corrected p-value (q-value) ≤ 0.01. All pathways passing these criteria were plotted. The species contributing these pathways were extracted from the pathway abundance table and grouped at the genus level. The proportion of each genus contributing each pathway was determined, and genera contributing < 5% were grouped together as “Other”. Plots were assembled using the R package patchwork ^19^.

All steps for HUMAnN3 functional analysis can be found in the R markdown files on the github page 02-scripts.backup/Cameroon_humann2_decontam.Rmd, 02- scripts.backup/Cameroon_humann2_heatmaps.Rmd, 2-scripts.backup/Cameroon_humann2_fa.Rmd, 02-scripts.backup/Cameroon_humann2_fxn.Rmd, 02-scripts.backup/Cameroon_humann2_species.Rmd, which use tidyverse formatting. HUMAnN2 was run with default settings.

### Taxonomic profiles with and without metagenome-assembled metagenomes

To determine if there are taxa present in the Baka and Nzime samples that are not represented in the NCBI RefSeq database, we used Kraken2 ^20^ and built two databases, one containing only RefSeq genomes (here, RefSeq_kraken) and the other containing those same RefSeq genomes and the metagenome-assembled genomes published by Pasolli et al. ^21^ (here, RefSeq_Pasolli_kraken). We used Kraken rather than MALT for this because MALT databases require NCBI taxonomic IDs for LCA analysis, which not all of the MAGs from Passoli, et al. have. For creating a custom database for Kraken2, we first retrieved the list of all available genomes on NCBI RefSeq (date of access: 17/10/2019) and then downloaded all genomes belonging to the kingdoms bacteria, archaea, and fungi as well all viral genomes. When more than ten genomes were available for a given taxon, we restricted the number of genomes for this taxon by randomly selecting ten genomes with an assembly level higher than “Contig” (seed for random sampling: 0).

Metagenome-assembled genomes (MAGs) from Pasolli et al. ^21^ were downloaded from the authors’ website (http://segatalab.cibio.unitn.it/data/Pasolli_et_al.html) and were added to the NCBI RefSeq genomes for the RefSeq_Pasolli database. For all MAGs that did not have an exact species match in NCBI, and therefore lacked an NCBI taxonomic ID, the taxonomic ID of the closest genome with a NCBI taxonomic ID was selected to represent this MAG. The Kraken2 database was built with Kraken v2.0 with default parameters. The files with the number of classified and unclassified reads were read into R, where the differences between databases for each group were visualized with boxplots using ggplot.

We determined the relative abundance of each phylum in supragingival dental plaque samples by averaging the relative abundance presented in three studies included in the Human Oral Microbiome Database (https://www.homd.org/download#abundance), including V1-V3 amplicons and V3-V5 amplicons from ^22^, shotgun data from ^23^ and as-yet-unpublished data from Dewhirst (Dewhirst 35×9 data). All steps for analyses can be found in the R markdown files on the github page 02- scripts.backup/Cameroon_kraken_RSPM_stats.Rmd, and 02- scripts.backup/Cameroon_kraken_RSPM_taxonomy.Rmd.

### Taxonomic ratios

All steps for analyses can be found in the R markdown files on the github page 02- scripts.backup/Cameroon_taxonomy_BF_ratio.Rmd.

## Supplemental Tables

**Supplemental table S2.** Dietary overlap between Baka and Nzime.

| Ethnic group | Dietary group | Item |
| --- | --- | --- |
| Baka and Nzime | Carbohydrate | cassava |
|  |  | plantain |
|  |  | rice |
|  | Vegetable | cassava leaves |
|  |  | cocoyam leaves |
|  |  | koko leaves (*Gnetum africanum*) |
|  | Fruit | papaya |
|  | Fatty intake | peanuts (*Arachis hypogaea*) |
|  |  | bush mangoes (*Irvingia spp.*) |
|  |  | palm nuts |
|  |  | palm oil |
|  | Meat | bushmeat |
|  | Spices | chili |
|  |  | salt |
|  |  | stock cubes |
|  | Beverage | alcohol |
|  | Other | avocado |
| Baka | Carbohydrate | yam (*Dioscorea spp.*) |
|  | Vegetable | mushrooms (various) |
|  | Fatty intake | moabi seed oil (*Baillonella toxisperma*) |
|  | Meat | smoked fish |
|  |  | snails |
|  | Beverage | freshwater crustaceans |
|  | Beverage | soda |
|  | Other | grilled maize |
| Nzime | Insect | palm larvae |
| Nzime | Fruit | sweet plantain |
|  | Fatty intake | palm oil |
|  | Meat | oil (store bought) |
|  | Meat | pork |
|  | Spices | *Ricinodendron heudelotii* bark |
|  | Beverage | fermented palm sap |

## Supplemental Figures

**Figure S1.**
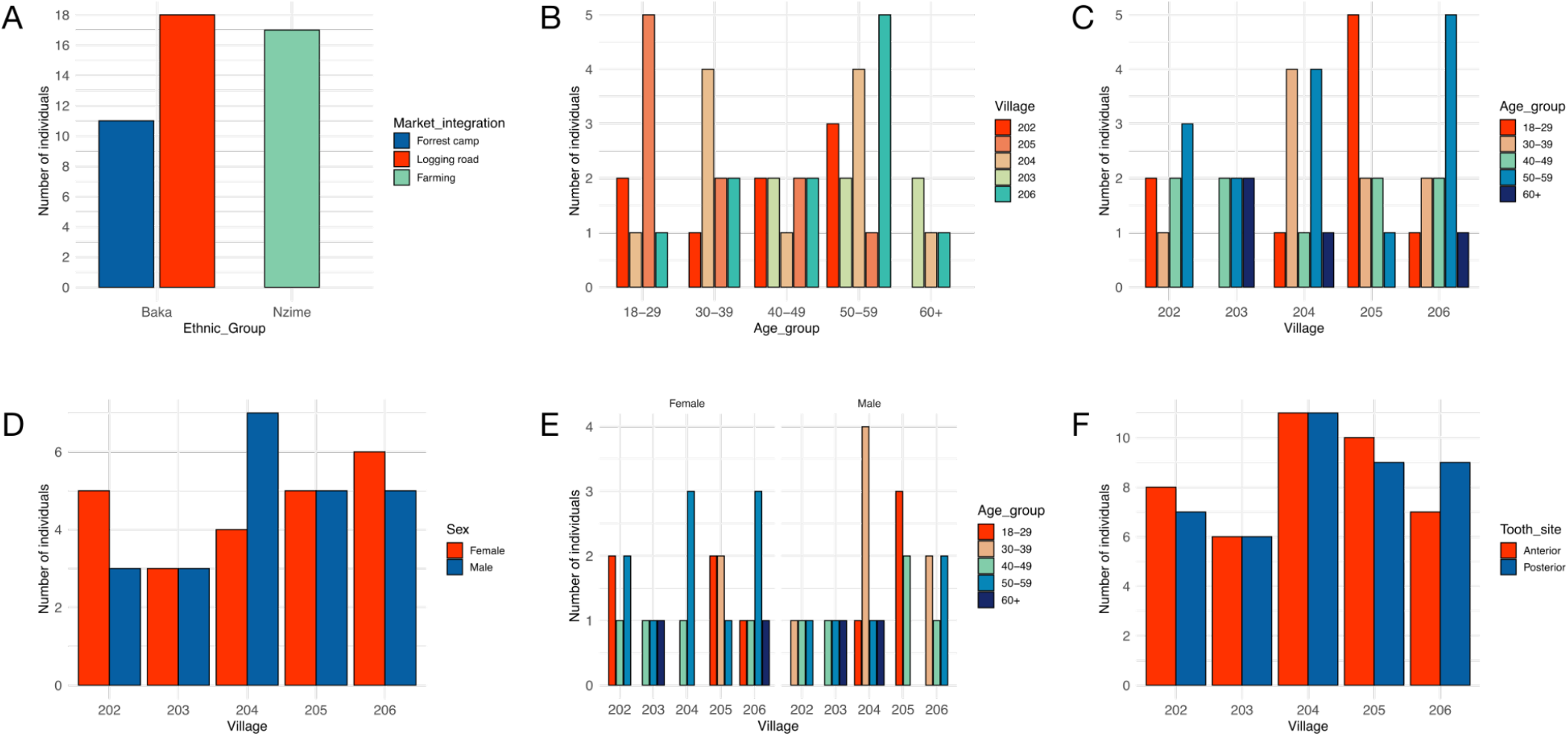
Participant demographics and sample information. The number of individuals broken down by **A**. market economy by ethnic group, **B**. village per age group, **C**. age group per village, **D**. sex per village, **E**. age group per village by male/female, **F**. tooth sampling site by village. Scripts are in the github page 02-scripts.backup/Cameroon_metadata_stats.Rmd.

**Figure S2.**
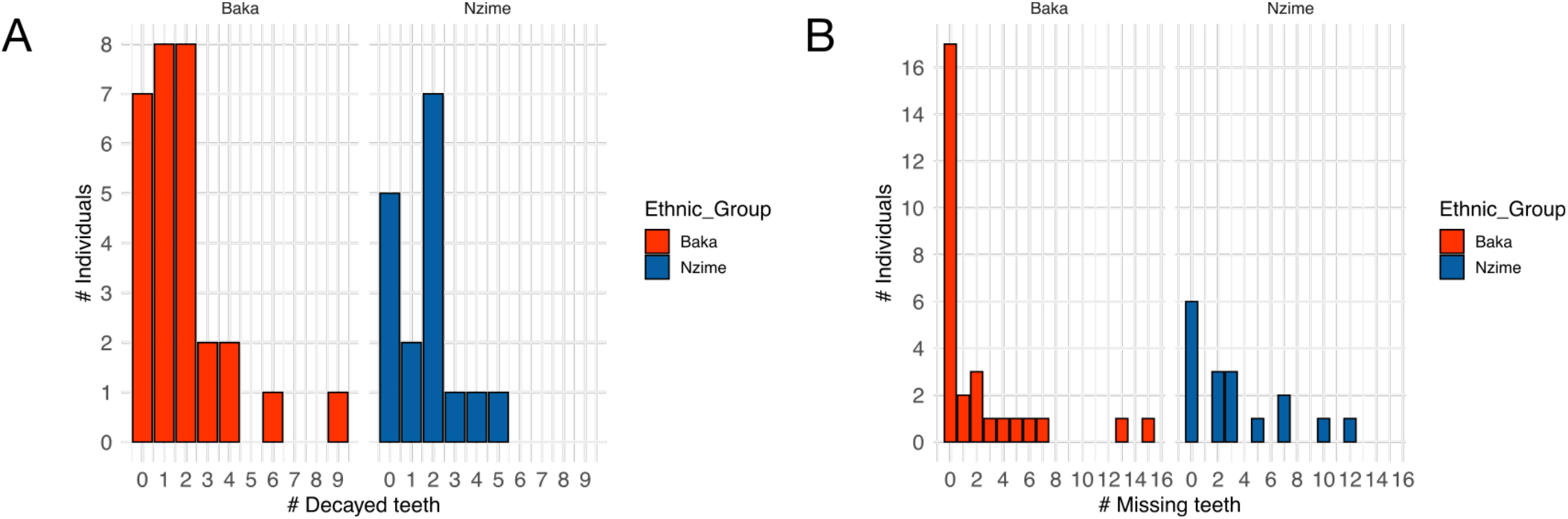
Decayed and missing teeth by individual. **A**. Number of individuals with 0, 1, or more decayed teeth, grouped by ethnic group. **B**. Number of individuals with 0, 1, or more missing teeth, grouped by ethnic group. Scripts are in the github page 02-scripts.backup/Cameroon_taxonomy_oral_health.Rmd.

**Figure S3.**
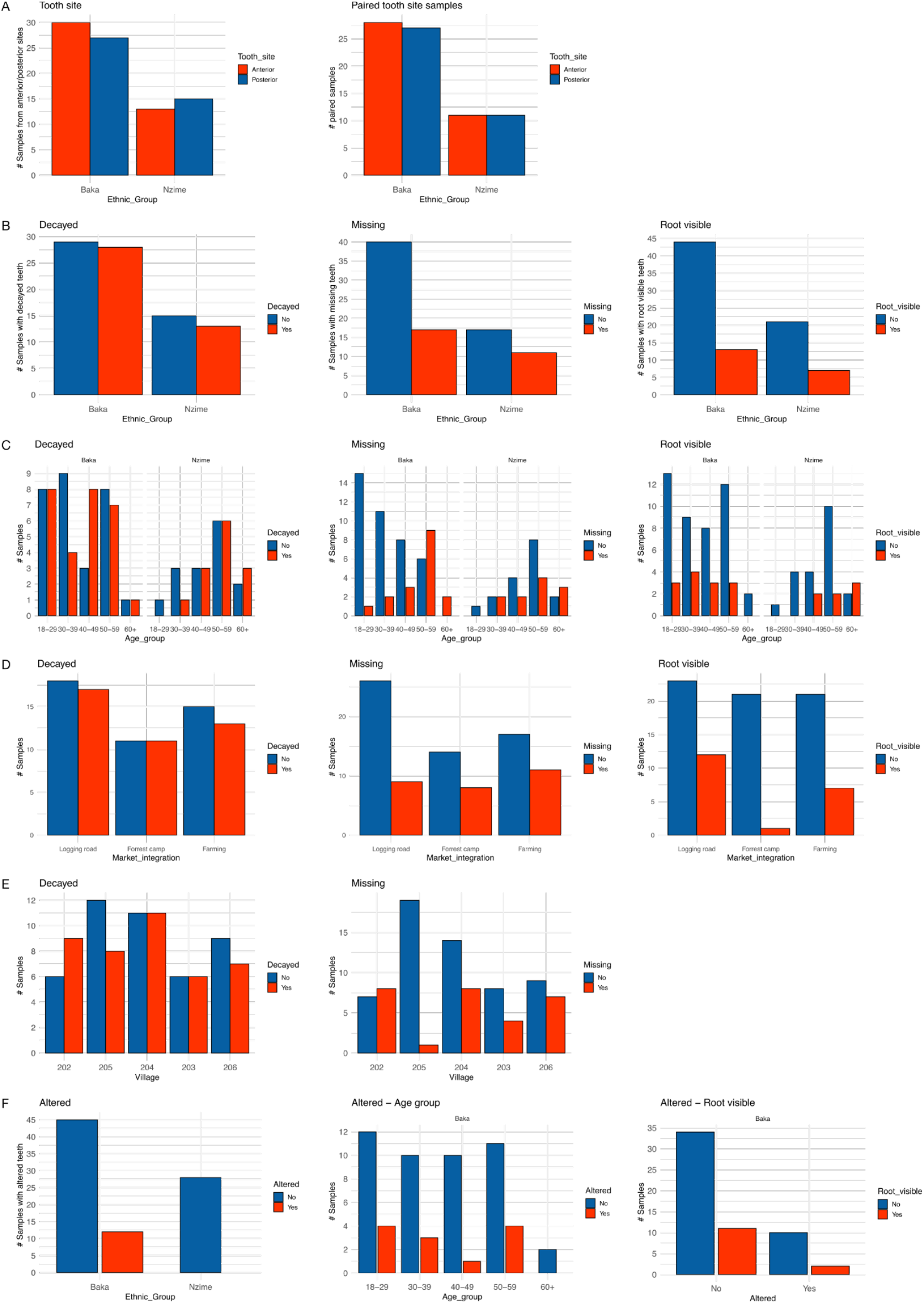
Oral health demographics. Distribution of samples **A**. from collection sites, having decayed, missing, or root-visible teeth **B**., by ethnic group **B**, age group **C**., market economy **D**., and village **D**. Tooth alteration **F**. predominantly of maxillary incisors, is practiced by some Baka, mostly women, but not by the Nzime. Scripts are in the github page 02- scripts.backup/Cameroon_taxonomy_oral_health.Rmd.

**Figure S4.**
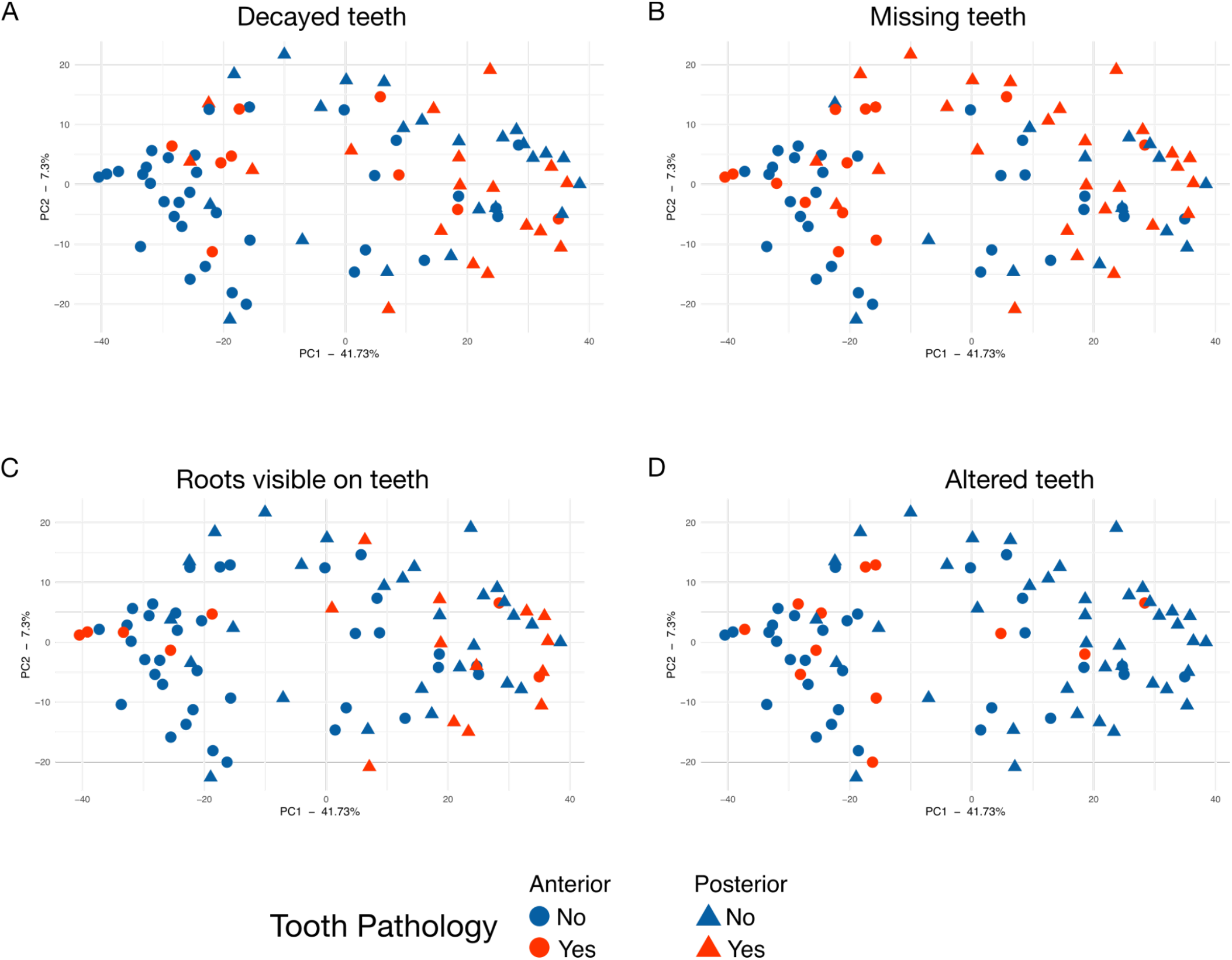
Pathology on teeth sampled from Baka and Nzime individuals does not cluster samples by species composition. PCA based on species composition with samples shaped by anterior or posterior collection sites, and colored by presence of **A**. Decayed teeth, **B**. Missing teeth, **C**. Teeth with visible roots, **D**. Altered teeth (mechanical filing). Only the Baka practice tooth alteration, which includes filing of the top anterior incisors. Scripts are in the github page 02- scripts.backup/Cameroon_taxonomy_oral_health.Rmd.

**Figure S5.**
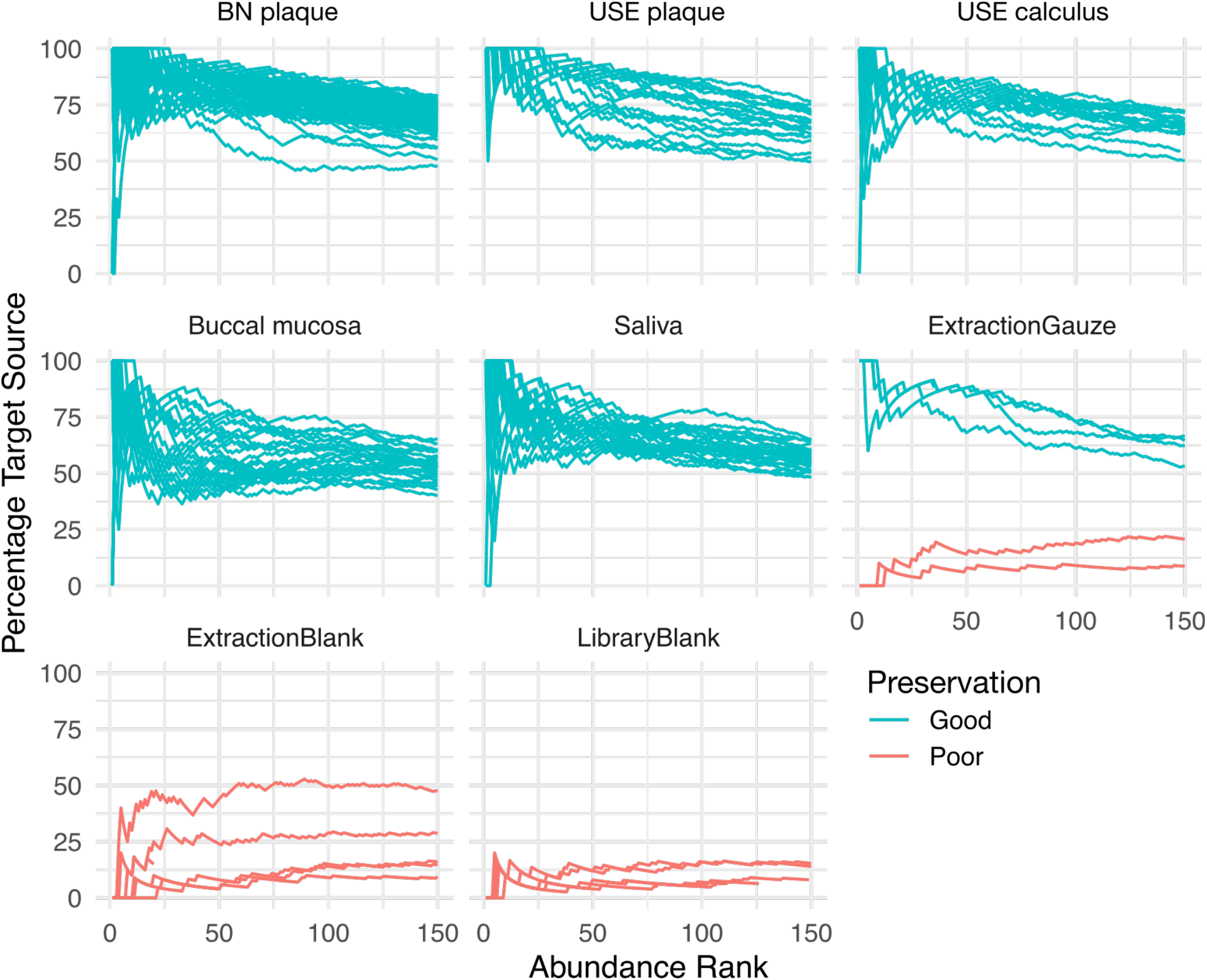
Cumulative percent decay curves. Adaptive burn-in filter with a threshold of 59%. Preservation - Good (turquoise lines) indicates the sample passes the threshold and is “well-preserved” with a sufficiently high proportion of oral taxa. Preservation - Poor (coral lines) indicates that the sample does not pass the threshold and is “poorly preserved”, with an insufficient proportion of oral taxa. The four extraction gauze samples that passed likely had saliva or small amounts of plaque that were not visible and were excluded from decontamination steps. Scripts are in the github page 02- scripts.backup/Cameroon_taxonomy_cuperdec.Rmd.

**Figure S6.**
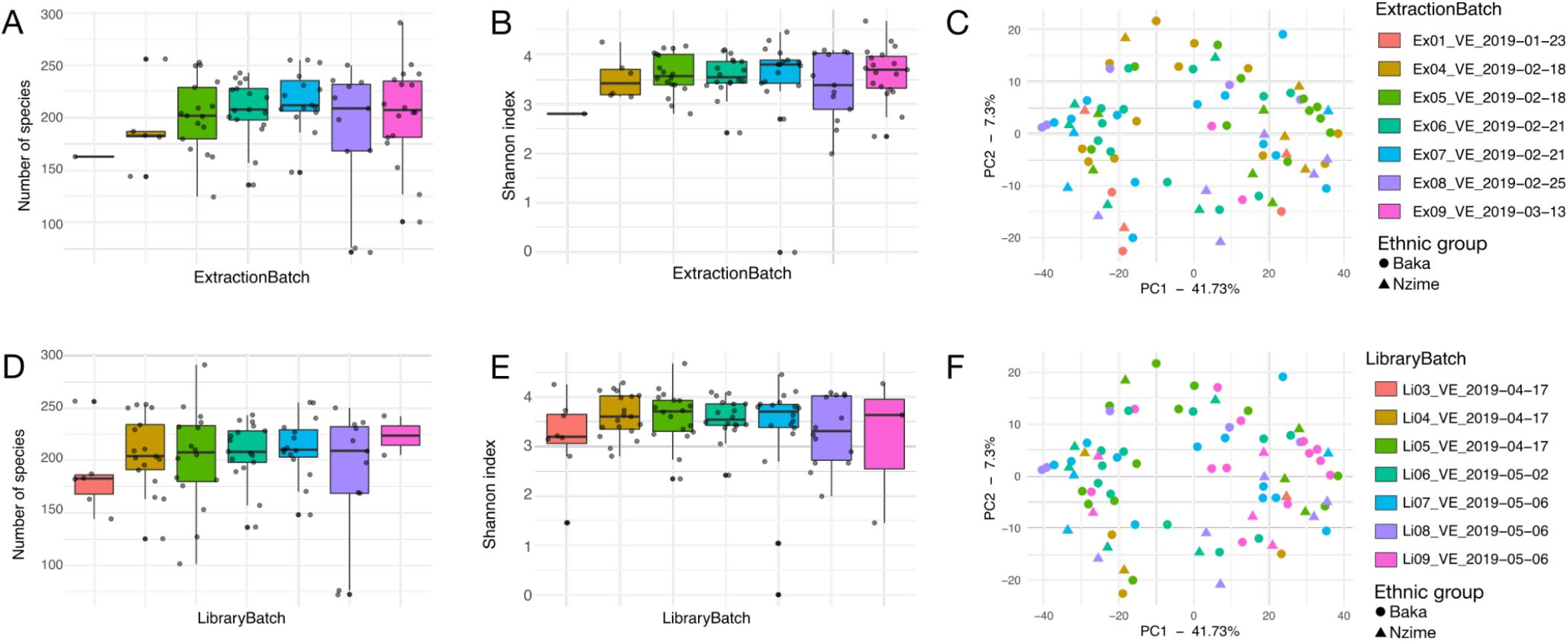
Microbial diversity of extraction batches. **A-C** Extraction batches. **A**. Number of species, **B**. Shannon index of species, **C**. PCA of species, **D-F** Library batches. **D**. Number of species, **E**. Shannon index of species, **F**. PCA of species. Statistical differences between alpha-diversity metrics (number of species/genera and Shannon index) were tested with Kruskal-Wallace test and Wilcox test with FDR correction. All statistical tests excluded EX01_VE_2019_01_23 because it had only a single sample. All test results had a p-value > 0.1. Statistical differences between groups in the PCAs were tested with PERMANOVA and all results had a p-value > 0.05. Alpha-diversity Scripts are in the github page 02- scripts.backup/Cameroon_taxonomy_batches.Rmd, and beta-diversity scripts are Cameroon_taxonomy_betadiv.Rmd.

**Figure S7.**
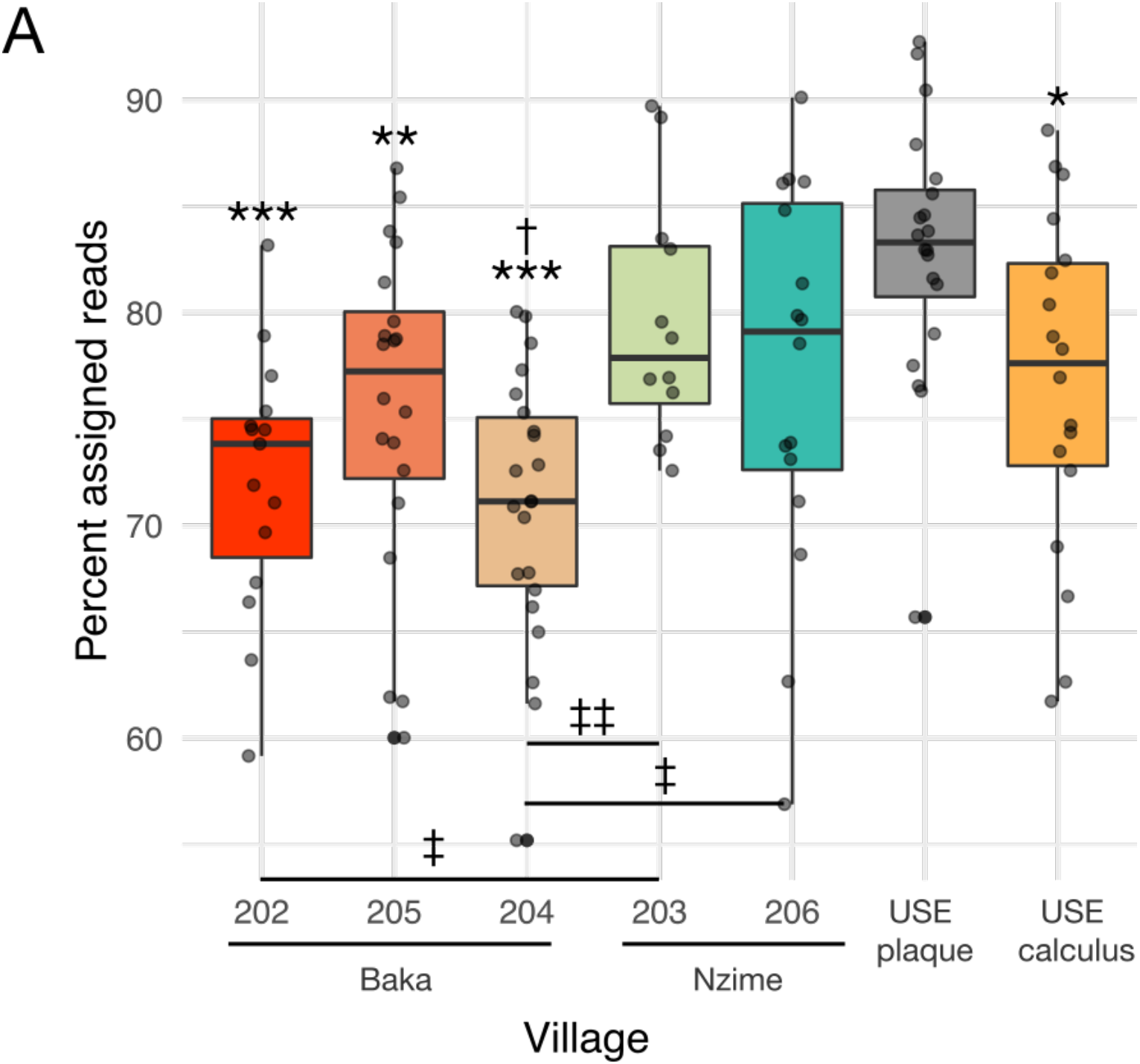
Percent of reads in each group that was assigned taxonomy by MALT. **A**. The number of reads assigned to BN plaque, USE plaque, and USE calculus samples with the NCBI RefSeq database. * p < 0.05, ** p < 0.01, *** p < 0.001 vs. USE plaque; † p < 0.05 vs. USE calculus; ‡ p < 0.05, ‡‡ p < 0.01 between villages. The script for this is found in Cameroon_plaque_assignment_stats.Rmd.

**Figure S8.**
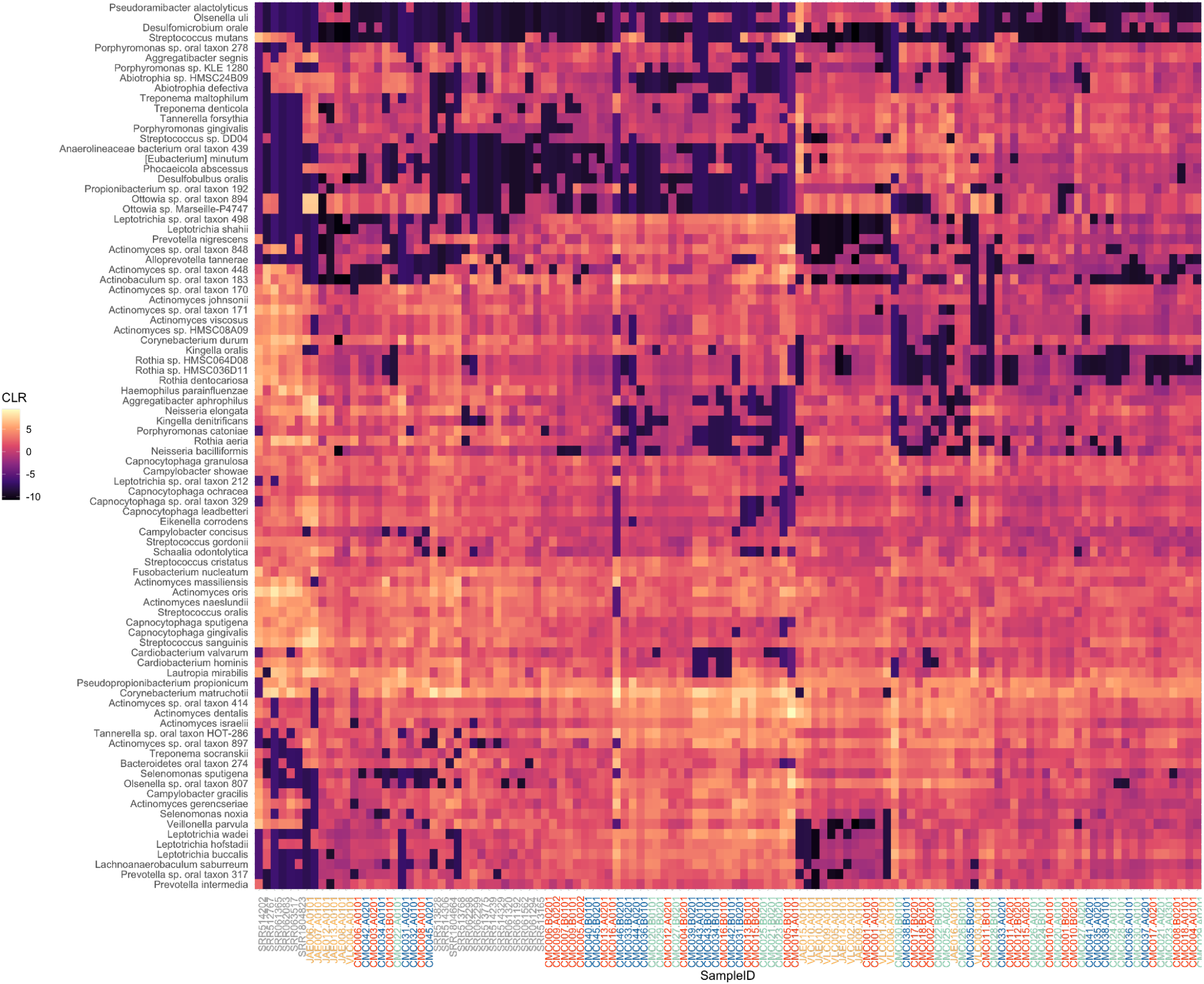
Heatmap showing the CLR-transformed abundance of the top 50 most abundant species in each group (BN plaque, USE calculus, USE plaque), clustered by both species and sample. Sample ID colors indicate market integration (red = Logging road (Baka), green = Forest camp (Baka), blue = Farming (Nzime), yellow = USE calculus, gray = USE plaque). Heat map scripts can be found in Cameroon_taxonomy_alphadiv.Rmd.

**Figure S9.**
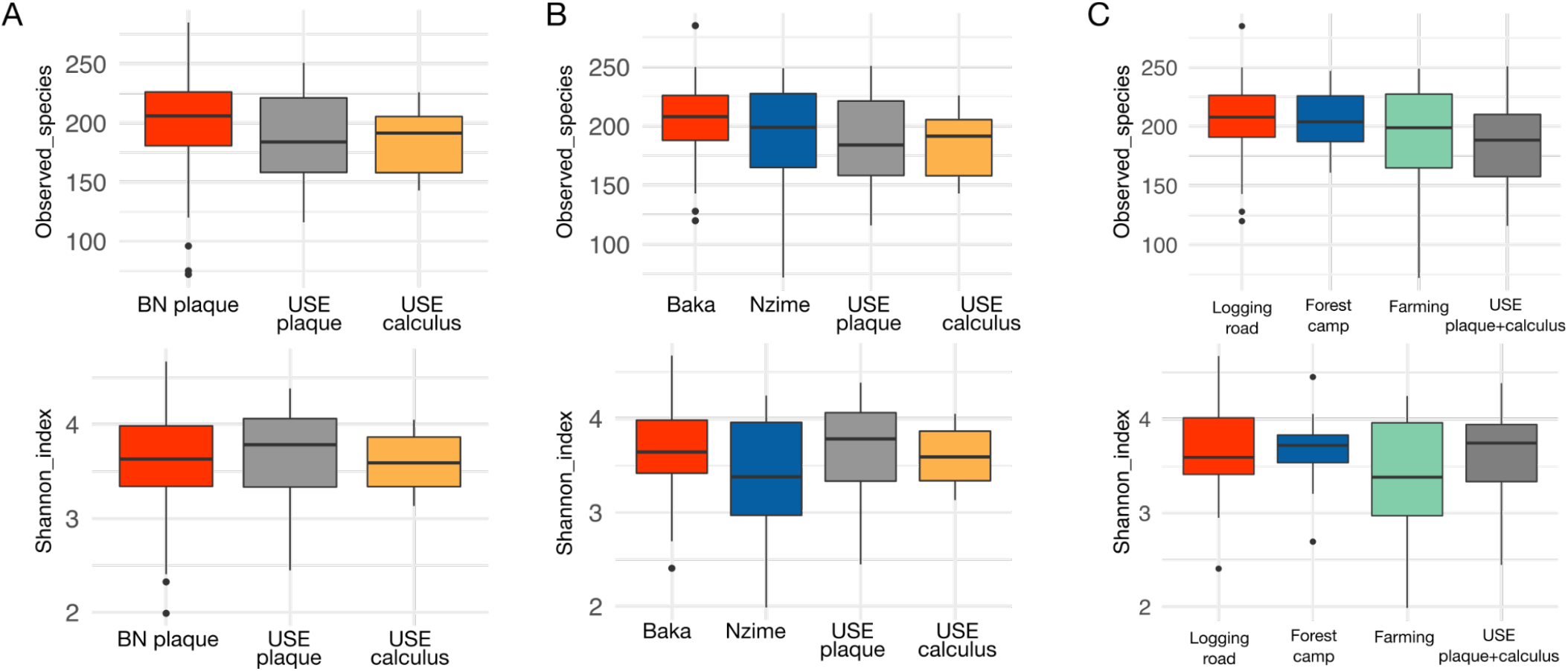
Species-level alpha-diversity metrics of all plaque and calculus sample groups. Indices include the number of species (top panel), and Shannon index (bottom panel). **A**. Sample source. **B**. Split by ethnic group and sample source. **C**. Market integration. No statistically significant differences within sample groups were found for any of the tested metrics. BN - Baka and Nzime; USE - US and European. Alpha-diversity scripts can be found in Cameroon_taxonomy_alphadiv.Rmd.

**Figure S10.**
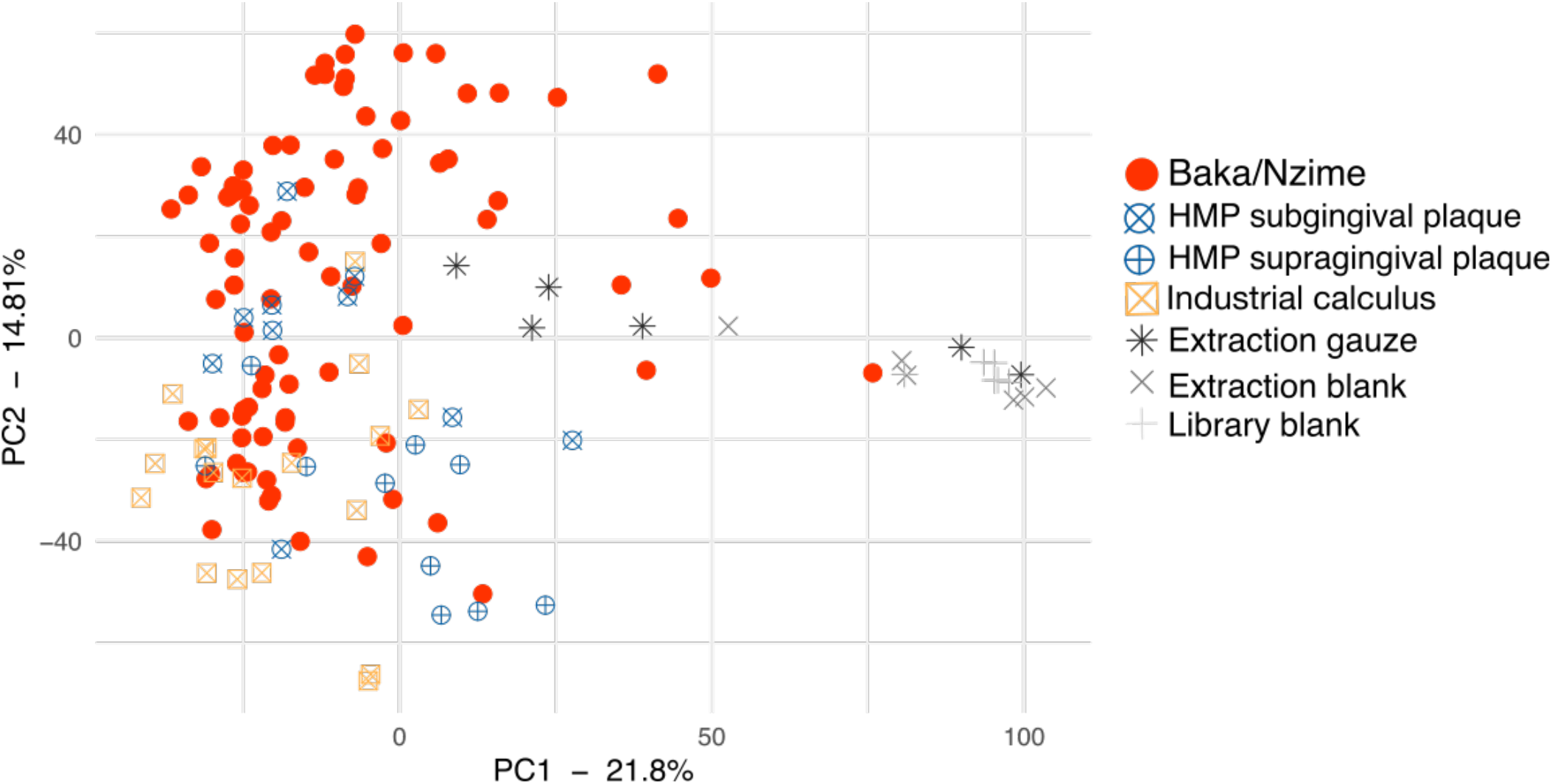
Raw species table PCA with all BN plaque, USE plaque, and USE calculus samples, as well as extraction and library blank samples. Species tables were not filtered for this analysis. The four extraction gauze samples that plot to the left of 50 on PC1 are the same four that passed the cuperdec threshold in Figure S5. Outlier samples were defined as those plotting to the right of 50 along PC1. These were removed in downstream analyses. The scripts for these plots are in Cameroon_taxonomy_betadiv.Rmd.

**Figure S11.**
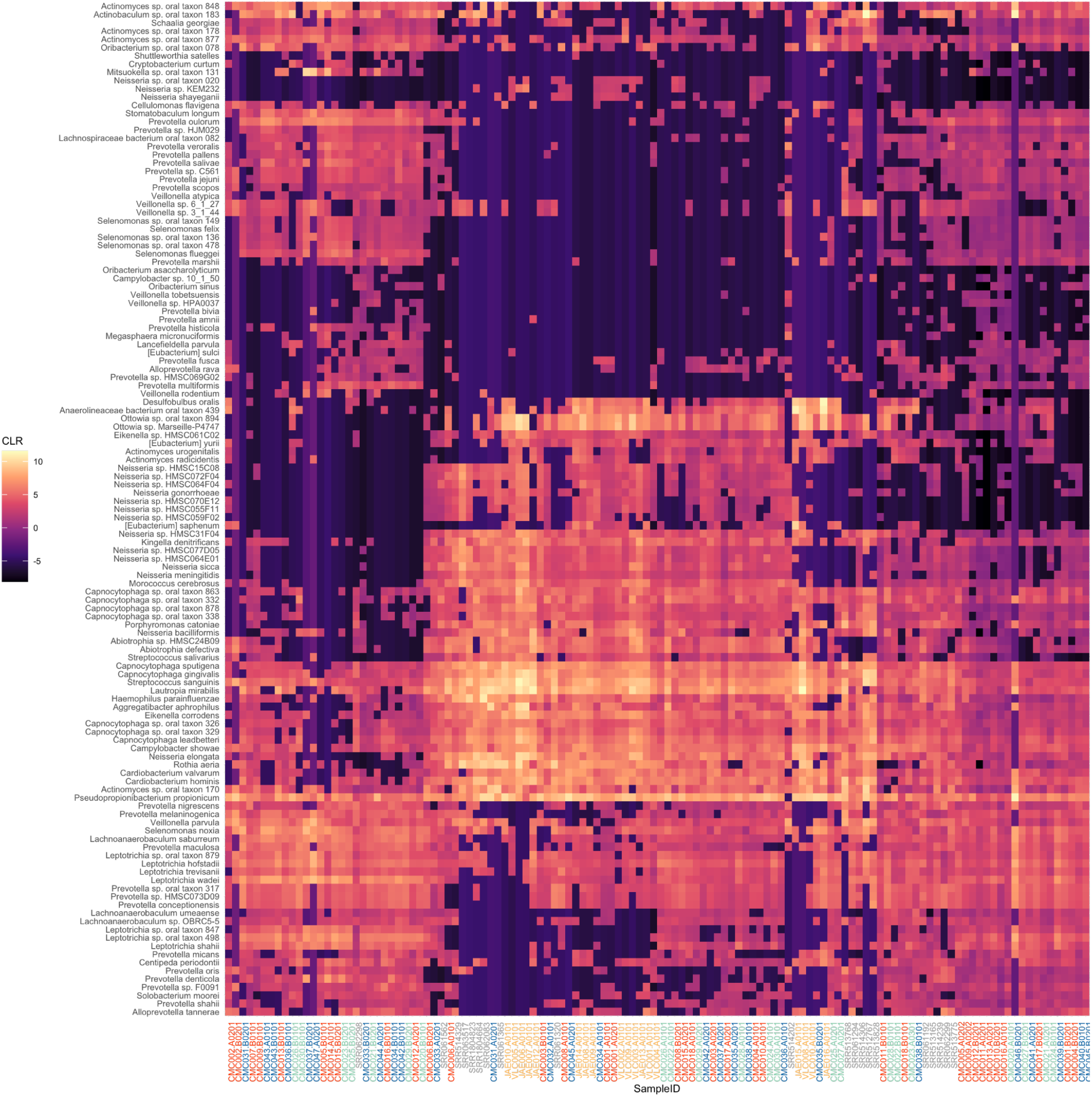
Heat map of the abundance of species in the first minimum residual from factor analysis of species in Baka and Nzime plaque, USE plaque and USE calculus samples. Values are CLR-transformed abundances. Sample ID colors indicate market integration (red = Logging road (Baka), green = Forest camp (Baka), blue = Farming (Nzime), yellow = USE calculus, gray = USE plaque). USE - US and European. All scripts can be found in Cameroon_taxonomy_discrimspecies.Rmd.

**Figure S12.**
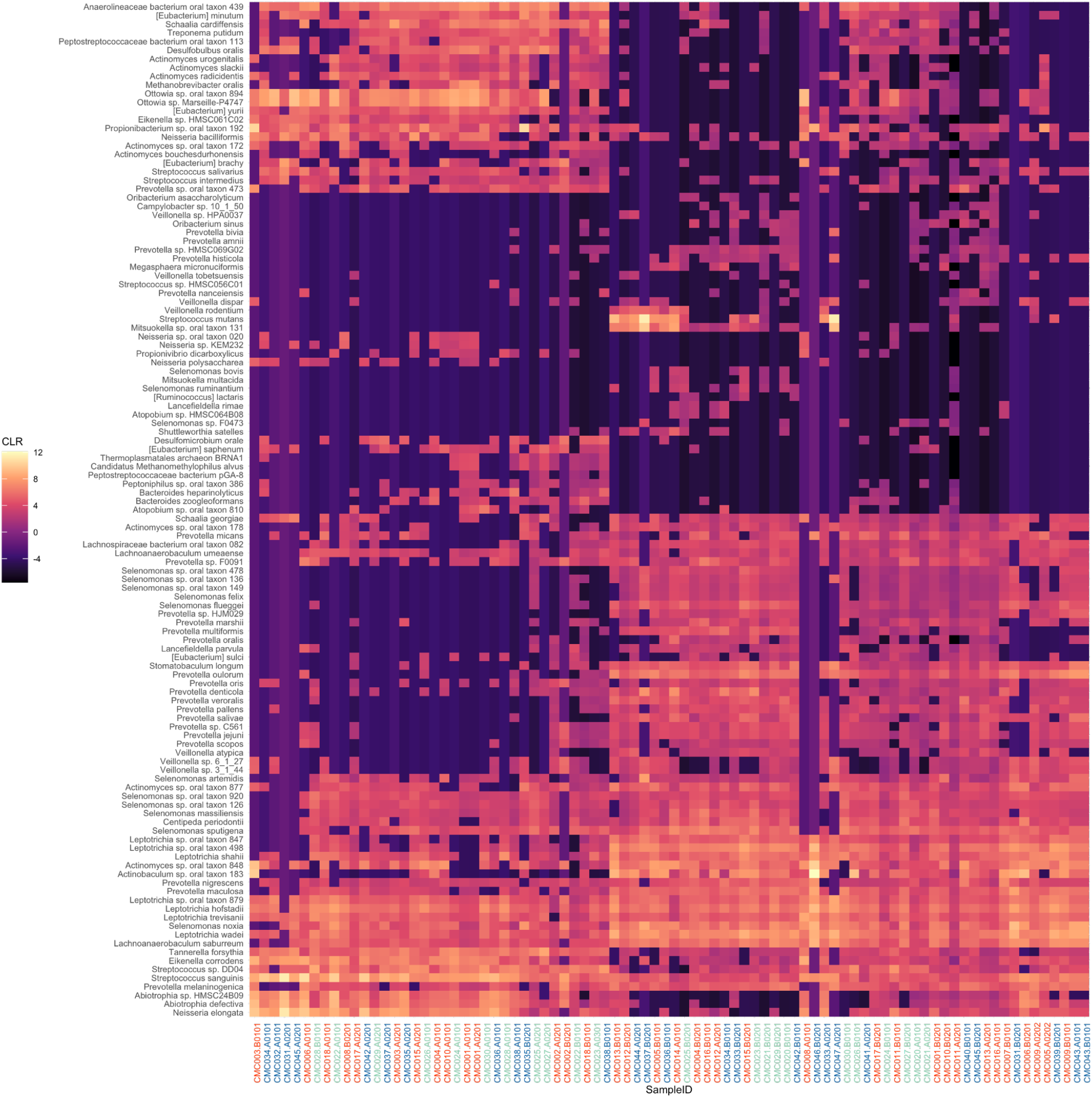
Heat map of the abundance of species in the first minimum residual from factor analysis of Baka and Nzime samples. Values are CLR-transformed abundances. Sample ID colors indicate market integration (red = Logging road(Baka), green = Forest camp (Baka), blue = Farming (Nzime)). All scripts can be found in Cameroon_taxonomy_discrimspecies.Rmd.

**Figure S13.**
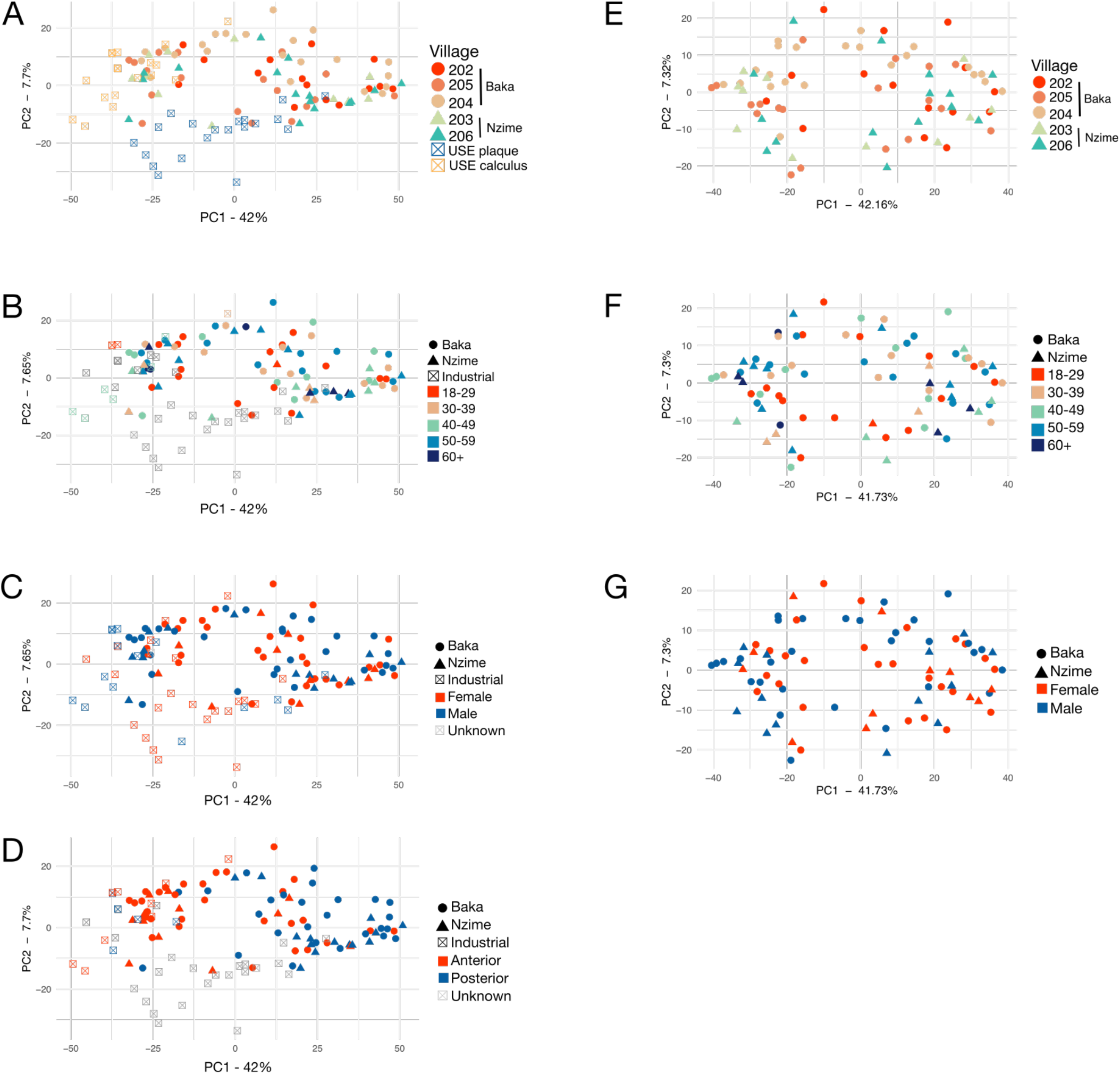
PCA plots based on species composition. **A-D** include USE plaque and calculus, **E-G** are Baka and Nzime plaque samples only. Plots are colored by **A**,**E**. Village, **B**,**F**. Age group, **C**,**G**. Sex., **D**. Tooth site. USE - US and European. All scripts including PERMANOVA and beta-dispersion tests can be found in Cameroon_taxonomy_betadiv.Rmd.

**Figure S14.**
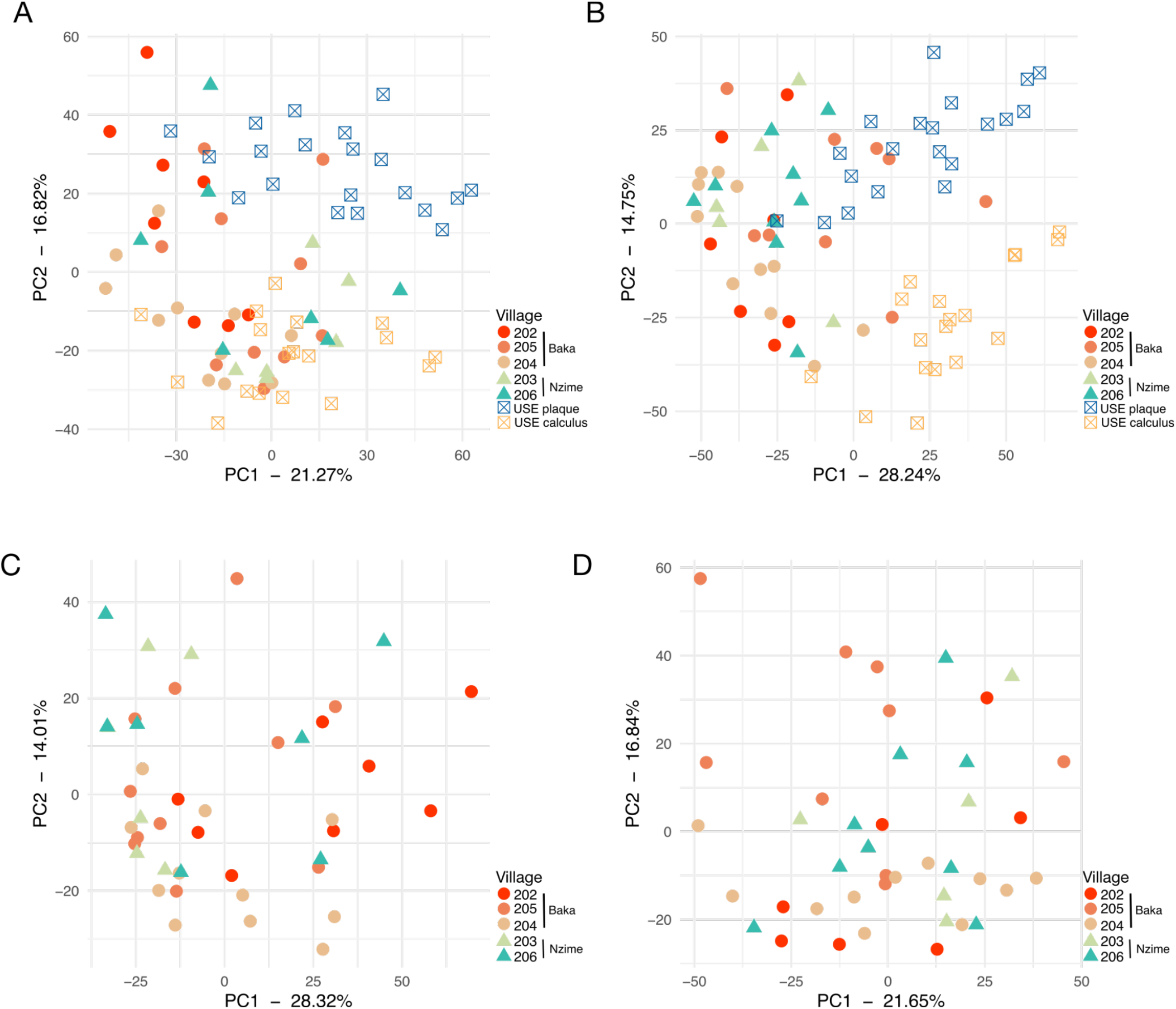
PCA plots based on species-level tables, with Cameroon plaque samples separately assessed by anterior (**A**,**C**) and posterior (**B**,**D**) tooth collection sites. Tables were filtered for species in the first 2 minimum residuals of factor analysis. USE - US and European. All scripts can be found in Cameroon_taxonomy_anterior_div.Rmd and Cameroon_taxonomy_posterior_div.Rmd.

**Figure S15.**
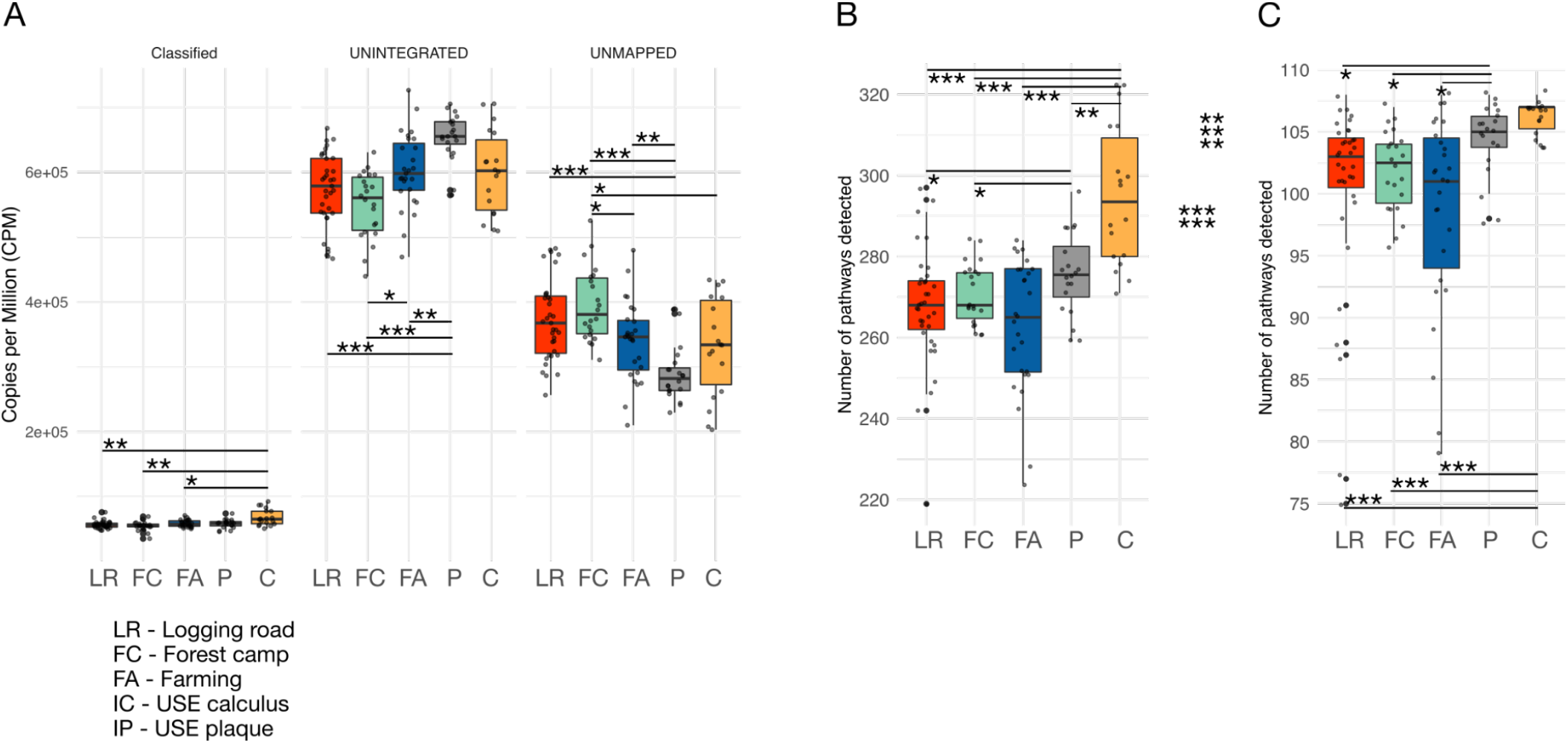
HUMAnN3 pathway analysis. **A**. Pathway abundance per million, which were classified, ungrouped, and unmapped by HUMAnN3, colored by level of market integration. **B**. Total number of pathways identified by HUMANn3 after removing potential contaminant pathways, colored by level of market integration. **C**. Number of pathways identified in the first minimum residual of factor analysis, colored by level of market integration. * p < 0.05, ** p < 0.01, *** p < 0.001. All scripts can be found in Cameroon_humann3_assignment_stats.Rmd and Cameroon_humann2_fxn.Rmd. USE - US and European; LR - Logging road; FC - Forest camp; FA - Farming; C - USE calculus; P - USE plaque.

**Figure S16.**
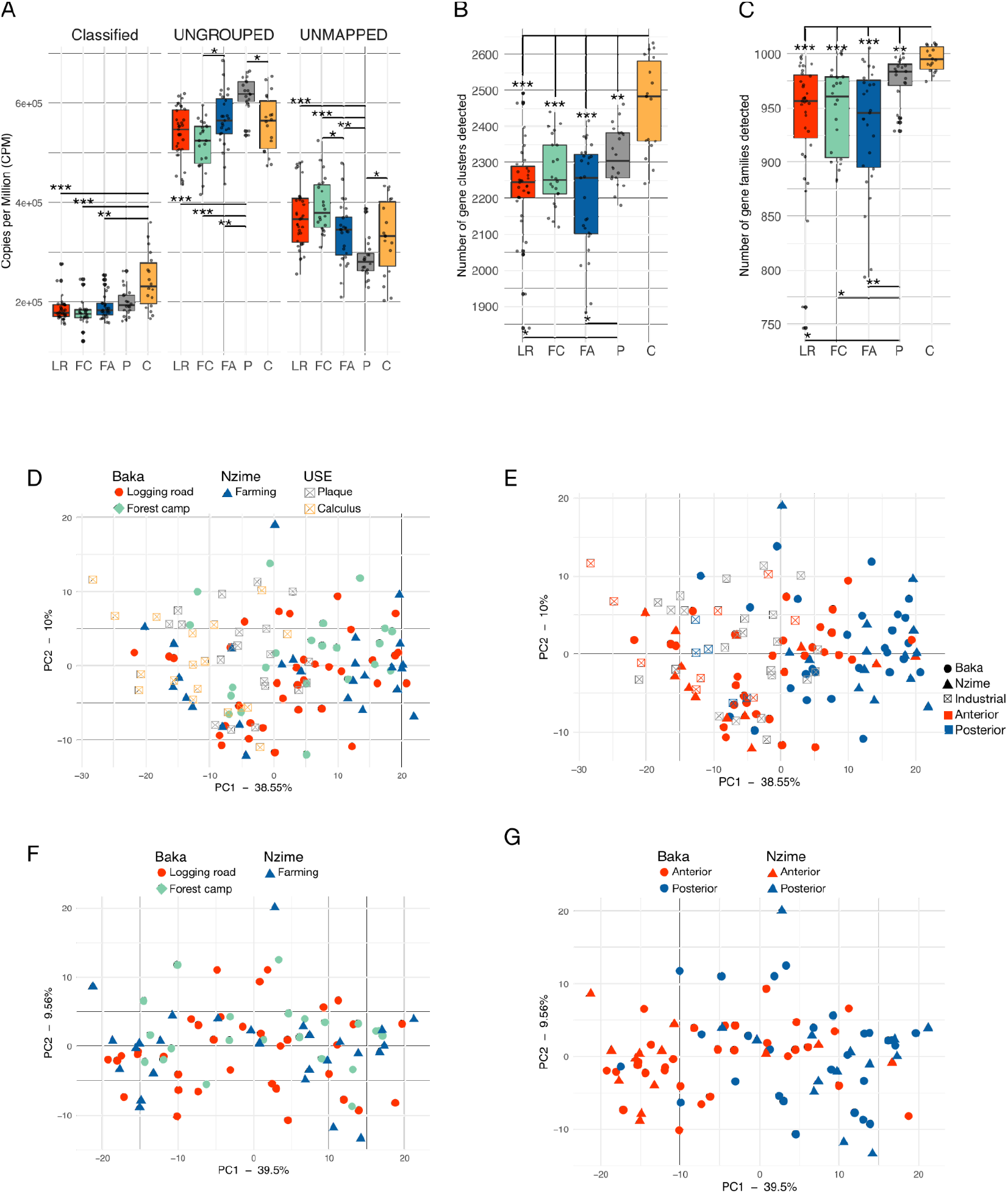
HUMAnN3 gene family analysis. **A**. Gene family copies per million that were classified, ungrouped, and unmapped by HUMAnN3, colored by level of market integration. **B**. Total number of gene clusters identified by HUMAnN3 after removing potential contaminant pathways. **C**. Number of gene clusters identified in the first minimum residual of factor analysis, colored by village. **D-E:** PCA plots based on gene clusters in the first minimum residual from factor analysis, including USE plaque and calculus, colored by **D**. level of market integration, **E**. tooth site. Gray indicates tooth site was not recorded, or plaque from multiple teeth were pooled. **F-G**: PCA plots based on gene clusters in the first factor from factor analysis, including only Baka and Nzime samples, colored by **F**. level of market integration, and **G**. tooth site. * p < 0.05, ** p < 0.01, *** p < 0.001. All scripts can be found in Cameroon_humann3_assignment_stats.Rmd and Cameroon_humann2_fxn.Rmd. USE - US and European. LR - Logging road; FC - Forest camp; FA - Farming; C - USE calculus; P - USE plaque.

